# Characterization of *aliA* and *aliB* deletion mutants reveals a dominant role of AliA in *Haloferax volcanii* lipoprotein lipidation

**DOI:** 10.64898/2026.03.08.710431

**Authors:** Yirui Hong, Andy Garcia, Samuel Kopelev, Jacob A. Cote, Friedhelm Pfeiffer, Paula Welander, Stefan Schulze, Mechthild Pohlschroder

**Author notes:** **Corresponding author:** Mechthild Pohlschroder; Stefan Schulze, **Email:**.

## Abstract

Protein lipidation is a widespread strategy for anchoring proteins to cellular membranes across all domains of life, yet the mechanisms underlying this process in archaea remain poorly understood. Recently, the first archaeal enzymes involved in lipobox-containing protein (lipoprotein) biogenesis, AliA and AliB, were identified and characterized in the model archaeon *Haloferax volcanii*. Although these paralogs share significant sequence similarity, distinct deletion phenotypes suggest differences in their substrate specificity and function. Here, we employed large-scale Triton X-114 fractionation followed by quantitative proteomics and lipid-specific mass spectrometry to systematically analyze AliA- and AliB-dependent lipoprotein lipidation. Deletion of *aliA* affected substantially more lipoproteins in *Hfx. volcanii* than deletion of *aliB*, markedly diminishing their TX-114 enrichment—indicating reduced hydrophobicity—and abolishing thioether-linked archaeol modification. This establishes AliA as the primary enzyme responsible for archaeal lipoprotein lipidation. In contrast, deletion of *aliB* affected only a small subset of lipoproteins and did not significantly reduce thioether-linked archaeol levels. In addition to defining distinct and non-redundant roles for AliA and AliB, this study provides the first large-scale experimental validation of predicted archaeal lipoproteins and identifies candidate components of the archaeal lipoprotein biogenesis pathway, substantially advancing mechanistic understanding and enabling improved lipoprotein prediction in this previously underexplored field.

## INTRODUCTION

Protein lipidation is a widely conserved phenomenon observed across all domains of life^1–4^. It significantly increases the hydrophobicity of proteins, thereby influencing their structures, stability, subcellular localization, enzymatic activities, and interactions with membranes and other proteins. Extensive studies in eukaryotes and bacteria have revealed a highly diverse landscape of lipidation types, each mediated by distinct enzymes and characterized by different conserved motifs, lipid anchors, biochemical properties, and biological functions^1–3,5^. In Archaea, a domain of life with important implications for eukaryogenesis^6–8^, ecology^9,10^, and biotechnology^11,12^, protein lipidation exhibits distinct features compared to the other two domains. While two lipidation types commonly seen in eukaryotes and/or bacteria, GPI anchoring^13^ and acylation^14–16^, have also been reported in archaea, the predominant form of archaeal protein lipidation is expected to be prenylation^17–24^, the addition of isoprenoid-derived moieties to proteins. This preference is likely attributed to the unique composition of archaeal membrane lipids, which are composed predominantly of isoprenoid-based alkyl chains ether-linked to glycerol-1-phosphate backbones^25–27^, in contrast to the fatty acyl chains ester-linked to glycerol-3-phosphate backbones in eukaryotes and most bacteria^28–30^.

Two major classes of prenylated proteins have been reported in archaea: the C-terminally lipidated ArtA (archaeasortase A) substrates^20,21,23,24,31^ and N-terminally lipidated proteins that contain a signature lipobox motif ([L/V/I]^−3^ [A/S/T/V/I]^−2^ [G/A/S]^−1^ [C]^+1^)^4,32^. These lipobox-containing proteins are also widespread in bacteria and are commonly referred to as bacterial lipoproteins in the literature^33,34^. In bacteria, lipoprotein biogenesis begins with the transport of lipoprotein precursors to cell membranes either in an unfolded state via the general secretory (Sec) pathway or in a folded state via the twin-arginine translocation (Tat) pathway^35^. The maturation of bacterial lipoproteins is subsequently carried out through a highly conserved three-step process. First, prolipoprotein diacylglyceryl transferase (Lgt) catalyzes the covalent attachment of a membrane lipid moiety to the lipobox cysteine^36^. Next, the lipoprotein signal peptidase (Lsp) cleaves the signal peptide N-terminal to the modified cysteine^37^. Finally, in some lipoproteins, the α-amino group of the cysteine undergoes further lipidation by lipoprotein N*-*acyltransferase (Lnt)^38^, lipoprotein *N*-acyl transferase system A and B (LnsAB)^39^, or lipoprotein intramolecular transacylase (Lit)^40^. A substantial fraction of archaeal proteins contain a canonical lipobox motif with an essential cysteine and are therefore considered to be archaeal lipoproteins^4,41^. However, despite this widespread genomic signature, only a small number of predicted archaeal lipoproteins have been experimentally confirmed^4,42–44^. Moreover, no homologs of the bacterial lipoprotein biosynthetic enzymes have been reported in archaea, consistent with the substantial differences in membrane lipid composition between the two domains.

Recently, by combining bioinformatic analyses with biochemical characterization, we identified two paralogous proteins, AliA and AliB, as the first known components of the archaeal lipoprotein biogenesis pathway^43^. While AliB is restricted to Haloarchaea, AliA is widely conserved across archaeal species. Deletion of *aliA* and/or *aliB* in the model archaeon *Haloferax volcanii* impaired lipoprotein biogenesis and caused diverse physiological defects, including slower growth, decreased motility, and altered cell shape. Deletion of both genes resulted in a complete loss of thioether-linked archaeol modification in lipoprotein extracts, demonstrating AliA and/or AliB are essential for archaeal protein prenylation. Notably, Δ*aliA* showed more severe defects than Δ*aliB* in both lipoprotein biogenesis and cell physiology, and the defect of one could not be complemented by overexpressing the other, indicating different enzymatic activities of the two paralogs. One possible explanation is that AliA serves as the primary enzyme for forming the thioether-linked archaeol, similar to the case of model bacterium *Mycobacterium smegmatis*, where one Lgt homolog is substantially more active than the other^45,46^. This is further supported by the observation that two conserved arginine residues in AliA correlate with Lgt arginine residues known to be critical for binding to negatively charged membrane lipids^47,48^, as revealed by visual structural comparison^43^, whereas these positions are replaced by two histidines in AliB^43^. Alternatively, AliA may act on a broader set of substrates than AliB, even if both possess comparable catalytic activity.

Here, we aimed to uncover mechanisms underlying the phenotypic differences between Δ*aliA* and Δ*aliB*. Using large-scale Triton X-114 extraction and subsequent mass spectrometry (MS), we systematically assessed the lipidation status of predicted lipoproteins in wild-type and mutant *Hfx. volcanii* strains. This approach not only resolved the distinct substrate specificities of AliA and AliB, but also substantially expanded the catalog of archaeal proteins experimentally validated as lipoproteins. In parallel, lipid-specific mass spectrometry revealed a marked reduction in thioether-linked archaeol modifications in Δ*aliA*, but not in Δ*aliB*. Together, these findings establish AliA as the dominant contributor to lipoprotein lipidation in *Hfx. volcanii*.

## RESULTS

### Triton X-114 extraction enriched hydrophobic proteins in *Hfx. volcanii*

To determine substrate specificities of AliA and AliB in *Hfx. volcanii*, we set out to examine the lipidation status of lipobox-containing proteins (i.e. predicted lipoproteins, hereafter referred to as lipoproteins) and identify those with impaired lipidation in Δ*aliA*, Δ*aliB*, and Δ*aliA*/Δ*aliB*. Previously, we successfully used Triton X-114 (TX-114), a detergent widely used for extracting hydrophobic proteins in eukaryotes^49–51^ and bacteria^52–54^, to demonstrate lipidation of two archaeal lipoproteins^43^. TX-114 effectively separated the lipidated proteoforms in the detergent phase (TX phase) from their non-lipidated counterparts in the aqueous phase (AQ phase). However, the efficiency of TX-114 for isolating archaeal lipoproteins in bulk has not been established so far. Therefore, we first performed TX-114 extraction in wild-type *Hfx. volcanii* (strain H53) and analyzed the proteome of the two phases via label-free, quantitative proteomics. About 64% of the 4222 predicted proteins encoded by *Hfx. volcanii* were quantified across the TX and AQ phases, a coverage in line with the most comprehensive single dataset in the Archaeal Proteome Project (ArcPP) (see Supplementary Fig.1 for a comparison of peptide and protein identifications)^55,56^.

**Supplementary Fig. 1.**
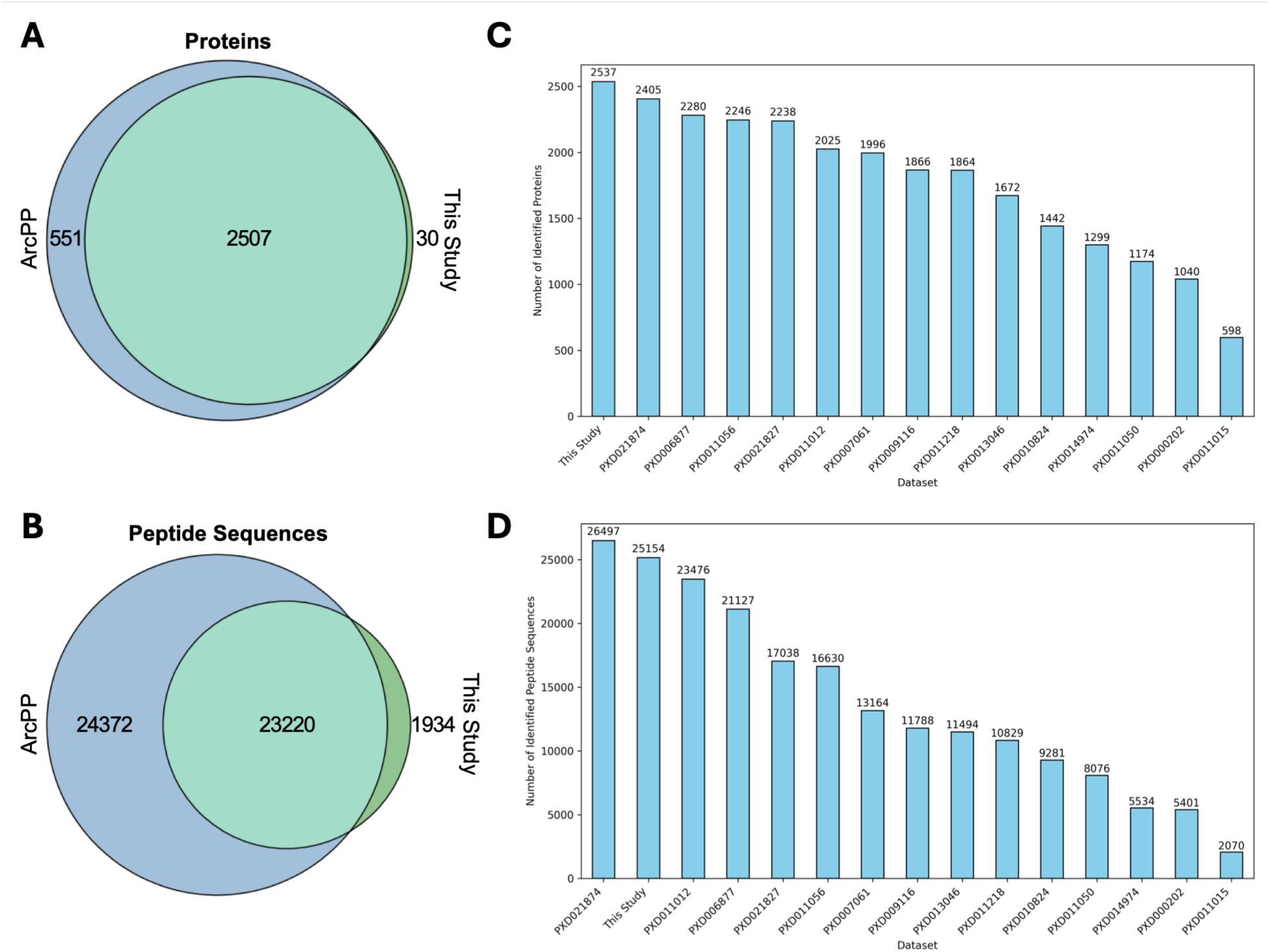
Proteome coverage comparison between this study and datasets within the Archaeal Proteome Project. Venn diagrams illustrating the overlap between **(a)** protein and **(b)** peptide identifications for this study and for the Archaeal Proteome Project (ArcPP, v. 1.4.0)^55^. Bar plots showing the number of identified **(c)** proteins and **(d)** peptides for this study and for each dataset in the ArcPP. The same filtering criteria were applied for all datasets (≤1% PEP on the peptide spectrum match level (PSM), ≤1% FDR on the peptide level, ≤0.5% FDR on the protein level, and at least 2 PSMs for each level). Notably, out of the 30 proteins that were uniquely identified in this study, 18 are predicted to have ≥2 TM domains, and 2 are predicted to be processed by a prepilin peptidase. These results indicate that fractionation with Triton X-114 is beneficial for achieving high proteome coverage, including a high coverage of membrane proteins. The dataset with the second-highest number of protein identifications and highest number of peptide identifications (PXD021874), published later than the original ArcPP study, used subcellular fractionation (cytosol, membrane, and culture supernatant) and two different enzymes for protein digestion (trypsin and GluC)^56^. It should also be noted that an additional large-scale proteomics study^57^ was published while this manuscript was in preparation, achieving the highest protein and peptide coverage to date using a different protein extraction and mass spectrometry workflow; the implementation of this recent dataset into the ArcPP is pending.

**Supplementary Table 1.** Protein counts by predicted group and status in the mass spectrometry analysis of detergent (TX) and aqueous (AQ) phase in wild-type *Hfx. volcanii* (strain H53). The numbers of proteins that were quantified only in the TX or AQ phase, or enriched in either phase, or showing no significant difference between the phases are listed. “Total” corresponds to the numbers of proteins for each predicted group within the whole theoretical proteome. ≥2 TM, proteins with at least two predicted transmembrane domains; Cyt, cytoplasmic proteins; Lipo, lipoproteins.

| Category | Total | Quantified | Only in TX | Enriched in TX | Only in AQ | Enriched in AQ | No significant difference |
| --- | --- | --- | --- | --- | --- | --- | --- |
| **≥2 TM** | 784 | 384 | 147 | 187 | 5 | 5 | 40 |
| **Cyt** | 3112 | 2045 | 22 | 397 | 107 | 1190 | 326 |
| **Lipo** | 108 | 87 | 13 | 59 | 0 | 2 | 13 |

We next examined the distribution of specific protein groups in the TX and AQ phases by comparing their relative abundances. As a positive control, proteins with two or more predicted transmembrane domains (hereafter referred to as ≥2 TM) were highly enriched in the TX phase (median log_2_FC of 3.71, Fig. 1a,b). In contrast, the vast majority of predicted cytoplasmic proteins (hereafter referred to as Cyt, defined as proteins lacking predicted transmembrane domains and secretion signals), as a negative control, were more abundant in the AQ phase (median log_2_FC of -1.1, Fig. 1a,b). Consistent with the overall high proteome coverage, 81% (87) of the 108 lipoproteins predicted by SignalP 6.0 with ≥90% confidence (Supplementary Data 1) were quantified and were highly enriched in the TX phase, in line with the anticipated behavior. Specifically, 72 (83%) of the 87 quantified lipoproteins were more abundant in the TX phase (Fig. 1a), of which 13 were detected exclusively in the TX phase while 59 were strongly enriched (Supplementary Table 1). Among lipoproteins showing significant abundance differences between the two phases, the median log_2_FC was 4.93 (Fig. 1b), which was significantly higher (p-value < 10^-10^) than that of Cyt proteins (log_2_FC = - 1.10) and even slightly higher (p-value = 0.055) than that of ≥2 TM proteins (log_2_FC = 3.71). Together, these results establish TX-114 extraction as an effective method to enrich hydrophobic proteins, providing a foundation for its application in analyzing lipoproteins across strains.

**Fig. 1.**
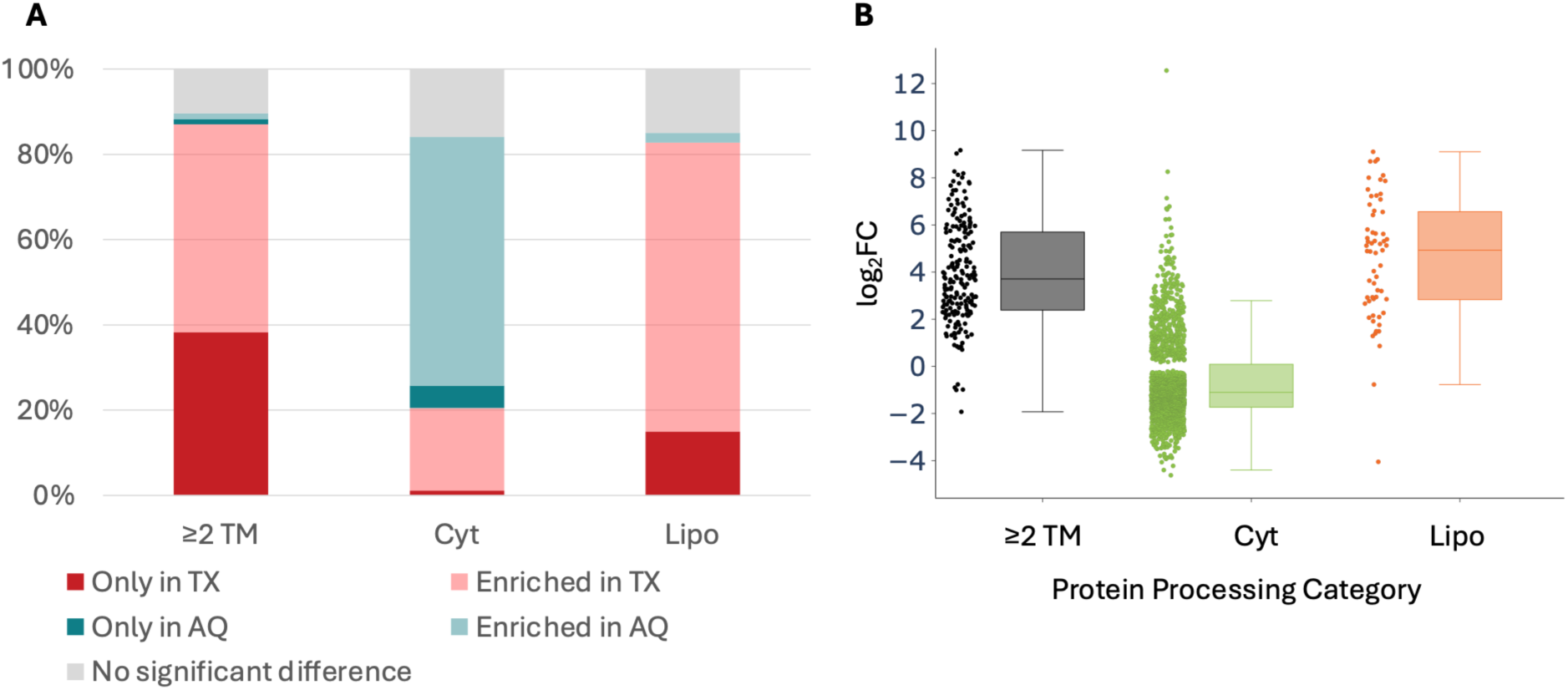
TX-114 efficiently enriched hydrophobic proteins in wild-type *Hfx. volcanii* (strain H53). **(a)** The stacked bar graph indicates the percentage of proteins that were quantified only in the TX-114 (TX) or aqueous (AQ) phase, or enriched in either phase, or showing no significant difference between the phases. Percentages for each predicted protein group are normalized to the total number of quantified proteins within that group. ≥2 TM, proteins with at least two predicted transmembrane domains; Cyt, cytoplasmic proteins; Lipo, lipoproteins. **(b)** Log_2_ fold-changes (FC) of protein abundance in TX versus AQ phase in wild-type *Hfx. volcanii* for different predicted protein groups are depicted as boxplots. Only proteins with significant abundance differences between the two phases are represented in the figure. Boxplots show the median (center line between boxes), interquartile range (boxes), as well as the range of data points within 1.5x of the interquartile range (whiskers).

### *AliA* deletion significantly reduced overall lipoprotein hydrophobicity

We next analyzed the protein profiles in the TX phase and AQ phase of Δ*aliA*, Δ*aliB*, and *ΔaliA*/Δ*aliB*. Although the number of quantified lipoproteins was comparable across all strains (Fig. 2a, Supplementary Table 2), the fraction of lipoproteins enriched or only quantified in the TX phase dropped from 82% (72 of 88) and 83% (73 of 88) in the wild type and Δ*aliB*, respectively, to 44% (39 of 88) and 42% (37 of 88) in Δ*aliA*/Δ*aliB* and Δ*aliA*, respectively (Supplementary Table 2). Accordingly, lipoproteins in the *aliA* deletion strains were increasingly enriched in the AQ phase or showed no significant difference between the two phases (Fig. 2a, Supplementary Table 2). While around 40% of lipoproteins remained associated with the TX phase in *aliA* deletion strains, this distribution does not necessarily indicate lipidation, as non-lipidated lipoproteins may still associate with the TX phase via the hydrophobic domain (H-domain) of their uncleaved signal peptides. Indeed, the median log_2_FC of lipoproteins showing significant abundance differences between the two phases decreased from 4.93 in the wild type to 1.02 in Δ*aliA*/Δ*aliB* and 1.10 in Δ*aliA*, reflecting an approximately 16-fold reduction in TX phase enrichment relative to the AQ phase, and thus a significant decrease in lipoprotein hydrophobicity (Fig. 2b, p-values < 10^-10^ for both strains compared to the wild type). In contrast, Δ*aliB* showed no significant change in log_2_FC (5.34 versus 4.93 in wild type, p-value = 0.84, Fig. 2b), suggesting that AliB has a much narrower substrate range than AliA or may function in processes other than lipidation. It should also be noted that no significant change in the TX/AQ distribution was observed for proteins with ≥2 TM domains in any of the mutant strains, indicating that the reduced protein hydrophobicity in Δ*aliA* and Δ*aliA*/Δ*aliB* is specific to lipoproteins (Fig. 2b).

**Fig. 2.**
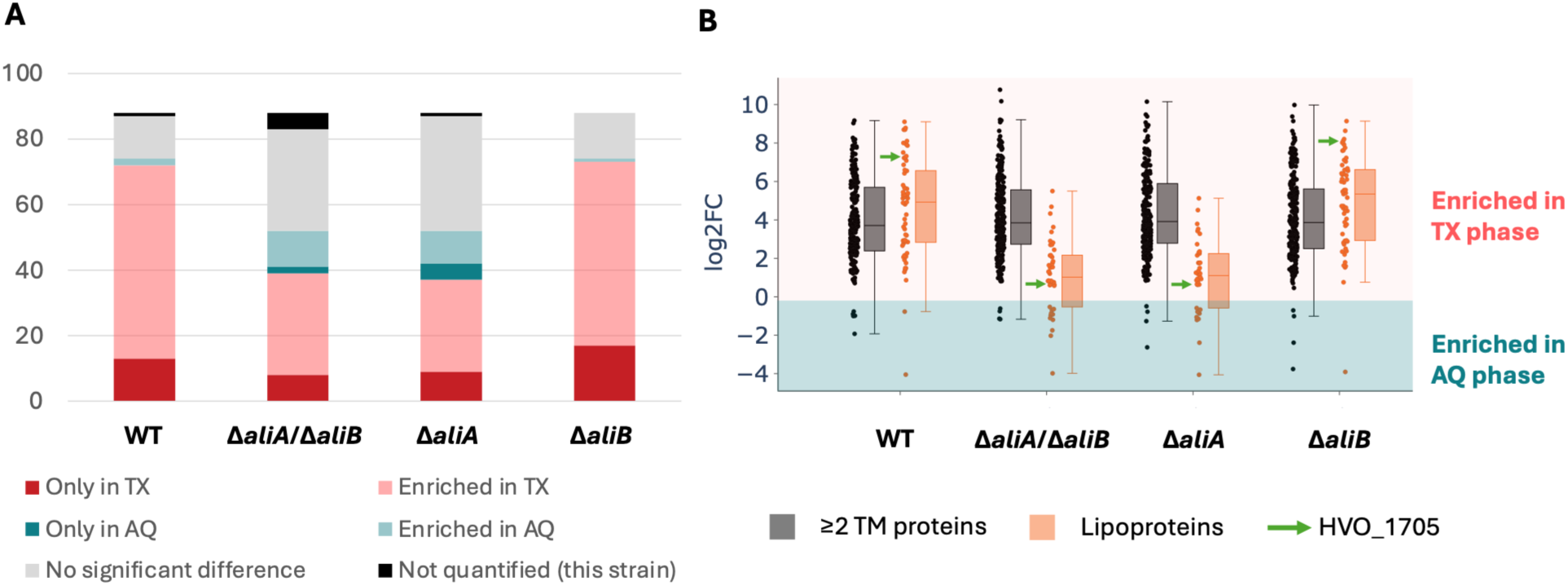
Deletion of *aliA*, but not *aliB*, significantly reduced the number of TX-enriched lipoproteins. **(a)** Number of lipoproteins quantified only in the TX-114 (TX) or aqueous (AQ) phase, or enriched in either phase, or showing no significant difference between the phases, across different strains. “Not quantified (this strain)” corresponds to the number of lipoproteins that have been quantified in other strains but not in the respective strain. **(b)** Log_2_ fold-changes (FC) of protein abundance in TX versus AQ phase in the analyzed strains for lipoproteins and ≥2 TM proteins (having at least two predicted transmembrane domains). Only proteins with significant abundance differences are represented in the figure. HVO_1705 is a verified Tat-transported lipoprotein^43^. Boxplots show the median (center line between boxes), interquartile range (boxes), as well as the range of data points within 1.5x of the interquartile range (whiskers).

**Supplementary Table 2.** Counts of *Hfx. volcanii* lipoproteins by enrichment status across strains. The numbers of proteins that were quantified only in the TX or AQ phase, or enriched in either phase, or showing no significant difference between the phases, or not quantified in the respective strain, are listed. “Total” refers to the number of lipoproteins quantified across all strains. “Not quantified (this strain)” refers to numbers of proteins that were detected in the whole MS dataset included in this study but were not quantified in the specific strain shown.

| Strains | Total | Quantified | Only in TX | Enriched in TX | Only in AQ | Enriched in AQ | No significant difference | Not quantified (this strain) |
| --- | --- | --- | --- | --- | --- | --- | --- | --- |
| **WT** | 88 | 87 | 13 | 59 | 0 | 2 | 13 | 1 |
| **ΔaliA/ΔaliB** | 88 | 83 | 8 | 31 | 2 | 11 | 31 | 5 |
| **ΔaliA** | 88 | 87 | 9 | 28 | 5 | 10 | 35 | 1 |
| **ΔaliB** | 88 | 88 | 17 | 56 | 0 | 1 | 14 | 0 |

### Large-scale proteomic validation of the predicted *Hfx. volcanii* lipoproteome

While several studies have predicted the prevalence and abundance of lipoproteins in archaea^4,41,43^, only a limited number of predicted lipoproteins have been experimentally validated^4,42–44^. To confirm additional lipoproteins, we applied a three-step validation strategy. First, we selected predicted lipoproteins that, in the wild-type sample, were significantly enriched in the TX phase compared to the AQ phase. From this subset, we identified proteins showing a significant reduction (at least 1.5-fold) of TX phase abundance in any of the three *ali* mutants compared to the wild type (defined as log2FC of TX_mut_:TX_WT_ < -0.58). Third, we excluded proteins that also exhibited reduced abundance in mutant AQ phase, thereby removing those non-lipoproteins with globally decreased expression in *ali* mutants and minimizing false positives. Together, this pattern was interpreted as indicative of reduced protein hydrophobicity in *ali* mutants and as supporting evidence for lipoprotein identity.

To validate this approach, we first assessed how previously verified lipoproteins performed in this analytical workflow. Four previously confirmed *Hfx. volcanii* lipoproteins met these criteria (Supplementary Data 2), supporting the robustness of the method. This included HVO_1245, which is predicted as cytoplasmic by SignalP 6.0 and would have been excluded based solely on prediction; however, our MS data supports its lipoprotein identity, highlighting the potential of this method for improving prediction tools. In line with the reliability of our approach, HVO_1808, a predicted Sec lipoprotein previously shown to be processed even when its lipobox cysteine was mutated^4^, did not pass the filter and displayed similar TX phase enrichment across all strains, suggesting its SignalP 6.0 lipoprotein prediction is likely a false positive (Supplementary Data 2).

Having validated the workflow, we applied it to all 88 predicted lipoproteins quantified across the dataset. After applying the three selection criteria, 51 proteins remained, supporting their predictions as lipoproteins (Fig. 3a). Most (40 out of 51) showed reduced hydrophobicity in strains lacking *aliA* (Δ*aliA* and Δ*aliA*/Δ*aliB*), but not in Δ*aliB* (Fig. 3a, Supplementary Table 3), indicating that AliB is involved in the lipidation of substantially fewer *Hfx. volcanii* lipoproteins than AliA. This group includes HVO_1705, a previously characterized lipoprotein that showed impaired lipidation in Δ*aliA* and Δ*aliA*/Δ*aliB* but remained largely unaffected in Δ*aliB*^43^ (Fig. 2B). In addition, five lipoproteins were uniquely affected in Δ*aliA* and three in Δ*aliA*/Δ*aliB* (Fig. 3a, Supplementary Table 3), suggesting minor differences between the two mutants. Of the three AliB substrates identified, one was affected only in *ΔaliB* (Fig. 3a, Supplementary Table 3).

**Fig. 3.**
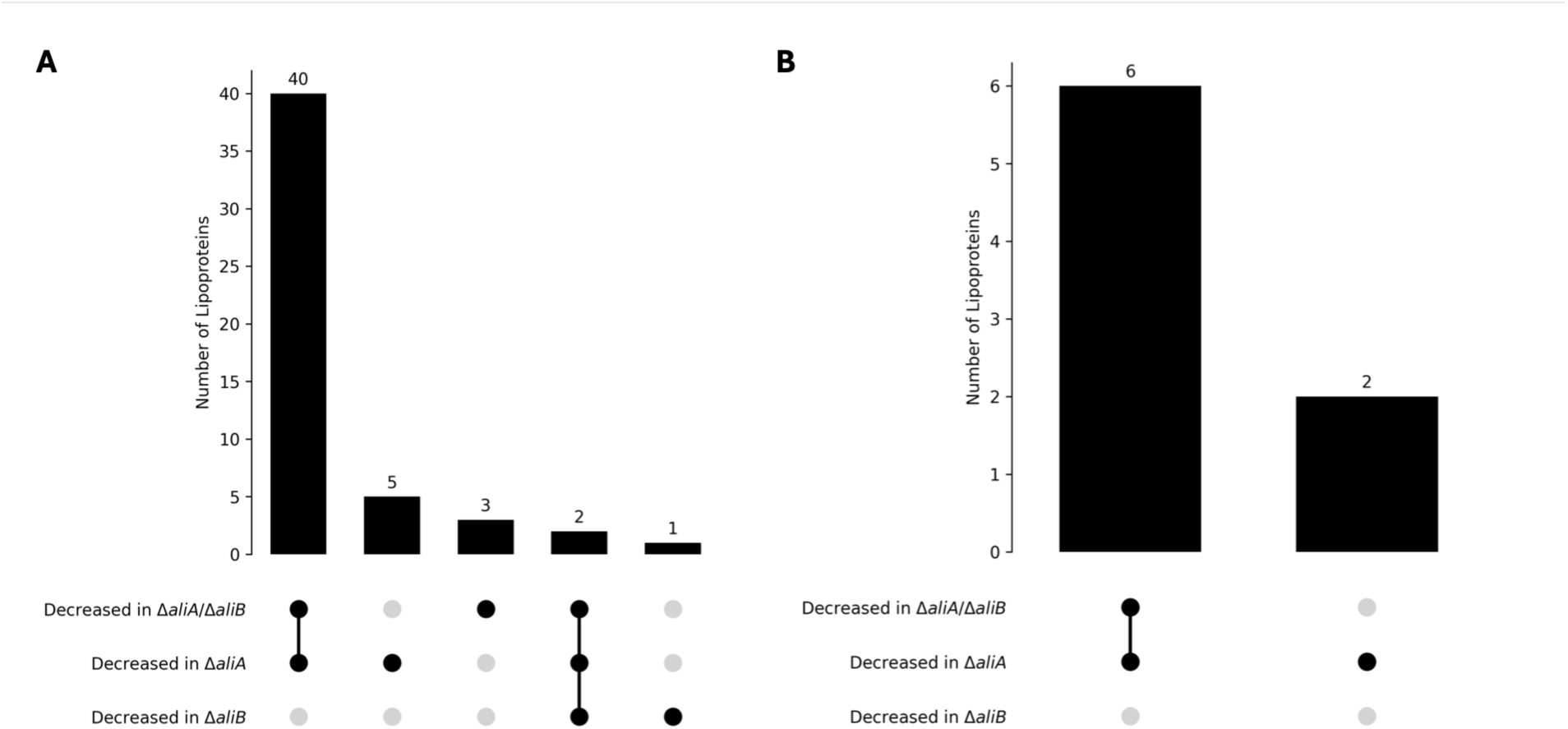
Large-scale identification of lipoproteins through their reduced TX-phase abundance in *ali* deletion strains. **(a)** Numbers of lipoproteins that were enriched in wild-type TX phase and exhibited at least a 1.5-fold reduction in TX phase abundance in Δ*aliA*/Δ*aliB*, Δ*aliA*, or Δ*aliB* compared to the wild type, but were not significantly decreased in the corresponding AQ phase. These proteins are considered validated lipoproteins. **(b)** Numbers of lipoproteins identified using the extended criteria, including those not quantified in the wild-type AQ phase or the mutant TX phase.

**Supplementary Table 3.** Lipoproteins validated through their reduced TX-phase abundance in *ali* deletion strains. Predicted lipoproteins were considered validated if they were enriched in the TX phase of wild type and showed at least a 1.5-fold reduction in TX phase abundance in Δ*aliA*/Δ*aliB*, Δ*aliA*, or Δ*aliB* compared to the wild type, but were not significantly decreased in the corresponding AQ phase. Proteins are grouped based on whether they were quantified in all relevant conditions for comparison (Fig. 3a) or showed missing values in either the wild-type AQ phase or the mutant TX phase (Fig. 3b). **\***HVO_B0034 was identified in both analyses but met the detection criteria only for Δ*aliA*/Δ*aliB* in the Fig. 3a analysis and only for Δ*aliA* in the Fig. 3b analysis.

| Proteins quantified in all relevant conditions (Fig. 3a) |  |  |
| --- | --- | --- |
| Category | Count | Proteins |
| Decreased in $\Delta aliA$ and $\Delta aliA/\Delta aliB$ | 40 | HVO_0062, HVO_0447, HVO_0530, HVO_0628, HVO_1007, HVO_1240, HVO_1241, HVO_1242, HVO_1244, HVO_1401, HVO_1477, HVO_1603, HVO_1705, HVO_1888, HVO_2094, HVO_2153, HVO_2316, HVO_2375, HVO_2432, HVO_2607, HVO_2695, HVO_2798, HVO_A0181, HVO_A0283, HVO_A0324, HVO_A0362, HVO_A0428, HVO_A0611, HVO_A0619, HVO_A0620, HVO_A0623, HVO_A0627, HVO_B0021, HVO_B0047, HVO_B0082, HVO_B0093, HVO_B0138, HVO_B0292, HVO_B0328, HVO_C0075 |
| Decreased in $\Delta aliA$ only | 5 | HVO_1673, HVO_1806, HVO_2651, HVO_B0106, HVO_B0198 |
| Decreased in $\Delta aliA/\Delta aliB$ only | 3 | HVO_B0034, HVO_B0139, HVO_B0318 |
| Decreased in $\Delta aliA/\Delta aliB$, $\Delta aliA$, and $\Delta aliB$ | 2 | HVO_0807, HVO_A0380 |
| Decreased in $\Delta aliB$ only | 1 | HVO_B0184 |
| Proteins missing in either WT AQ phase or the mutant TX phase (Fig. 3b) |  |  |
| Category | Count | Proteins |
| Decreased in $\Delta aliA$ and $\Delta aliA/\Delta aliB$ | 6 | HVO_1119, HVO_1144, HVO_2286, HVO_A0549, HVO_B0144, HVO_B0228 |
| Decreased in $\Delta aliA$ only | 2 | HVO_A0557, HVO_B0034* |

Because the three-step workflow required statistical comparisons between TX and AQ phases in the wild type and between TX phases of wild type and mutant strains, proteins had to be quantified in all relevant fractions to meet the significance criteria. Consequently, proteins with missing values in either the AQ phase of the wild type or the TX phase of the mutant could not be evaluated in the initial analysis. However, such missing values are themselves consistent with strong phase enrichment and therefore consistent with lipoprotein behavior. We therefore applied extended criteria to capture these cases, resulting in the identification of seven additional validated lipoproteins (Fig. 3b, Supplementary Table 3). In addition, one protein included in Fig. 3b had already been identified in Fig. 3a, as it was detected in Δ*aliA*/Δ*aliB* in the first analysis but met the extended detection criteria only for Δ*aliA* in this subset (Supplementary Table 3). Among the eight proteins, six exhibited reduced hydrophobicity in both Δ*aliA* and Δ*aliA*/Δ*aliB*, while two were affected only in Δ*aliA* (Fig. 3b, Supplementary Table 3). Together, these results significantly expand the set of validated *Hfx. volcanii* lipoproteins from fewer than ten to over fifty and further underscore the marked difference in lipidation capacity between AliA and AliB.

### Localization and abundance of the Tat lipoprotein HVO_1705 differ between Δ*aliA* and Δ*aliB*

Building on the global differences observed across the lipoproteome, we next focused on an individual lipoprotein to further explore the distinct roles of AliA and AliB. Current proteomic data indicate that deletion of *aliA*, but not *aliB*, significantly affects the lipidation of the lipoprotein HVO_1705 (Fig. 2B, Supplementary Table 3). This is consistent with previous Western blot analyses, which suggest that deletion of *aliB* has only a minor effect on HVO_1705 lipidation but a more pronounced effect on its signal peptide cleavage^43^. Together, these observations highlight HVO_1705 as a suitable model for dissecting AliA- and AliB-dependent steps in lipoprotein biogenesis. We therefore attempted to purify HVO_1705 for direct MS analysis of both peptides and lipid modifications. Although purification was limited by low protein abundance and purity, during the process, we observed pronounced differences in its cellular localization among strains. Cell fractionation of strains overexpressing HVO_1705 showed that the protein was predominantly membrane-associated in the wild type (Fig. 4a). In contrast, in both Δ*aliA*/Δ*aliB* and Δ*aliA* strains, HVO_1705 was primarily detected in the cytoplasmic fraction and exhibited reduced overall abundance (Fig. 4a,b). These observations suggest that *aliA* deletion either impairs membrane targeting of overexpressed HVO_1705 or triggers the clearance of non-lipidated HVO_1705 from both the membrane and cytoplasm via an unidentified quality-control mechanism. Conversely, in the Δ*aliB* strain, the majority of HVO_1705 remained membrane-associated, including both the mature form and the presumed lipidated but non-cleaved intermediate form previously reported^43^. This pattern further confirmed that AliB contributes more prominently to signal peptide cleavage than to lipidation of HVO_1705. As a control, we examined the Tat-translocated non-lipoprotein HVO_0844, which is membrane-anchored by a C-terminal transmembrane segment rather than covalent lipid modification. Its membrane localization was unaffected by deletion of *aliA* or *aliB* (Fig. 4c,d), confirming that the localization effect observed in the mutants is specific to lipoproteins. Collectively, these results further highlight the distinct roles of AliA and AliB and suggest that *aliA* deletion may additionally reduce lipoprotein abundance through a mechanism that remains to be elucidated.

**Fig. 4.**
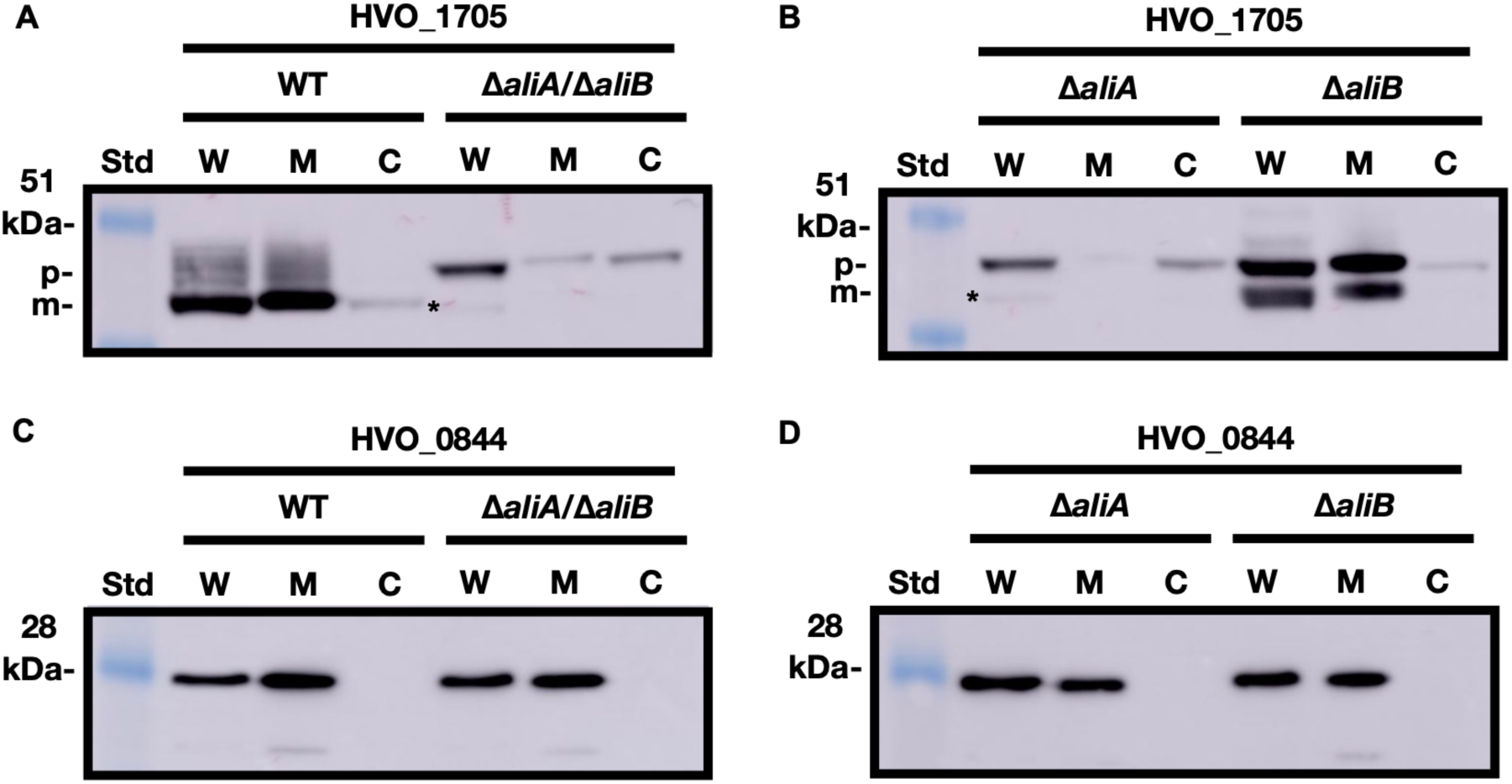
Deletion of *aliA*, but not *aliB*, led to a significantly reduced amount of membrane-associated HVO_1705, while the Tat-transported transmembrane protein HVO_0844 was unaffected by either deletion. Western blot of overexpressed myc-tagged HVO_1705 in **(a)** wild type, Δ*aliA*/Δ*aliB*, **(b)** Δ*aliA*, and Δ*aliB* in different cell fractions across strains. Western blot of overexpressed myc-tagged HVO_0844 in **(c)** wild type, Δ*aliA*/Δ*aliB*, (**d)** Δ*aliA*, and Δ*aliB* in different cell fractions across strains. Std, SeeBlue Plus2 pre-stained protein standard. W, whole cell lysates. M, membrane fraction. C, cytoplasmic fraction. Precursors (p) and mature proteins (m) are labeled accordingly. The asterisk indicates a likely degradation product of HVO_1705, as previously reported by Giménez et al^42^. The results are representative of two independent experiments. The original blots for panels a-d are included in the Source Data file.

### AliA is the main enzyme for thioether-linked archaeol modification of *Hfx. volcanii* lipoproteins

The results presented above focused on dissecting the differences between AliA and AliB using protein-based readouts. To complement these findings, we next examined their roles by directly analyzing lipid modifications across different strains. Previously, we showed that the TX-114 extract from the wild type contained methylthio-archaeol, the cleavage product of thioether-linked archaeol following methyl iodide treatment, whereas this modification was absent in Δ*aliA*/Δ*aliB*^43^. To determine which enzyme was responsible for this phenotype, we performed liquid chromatography-mass spectrometry (LC-MS) analysis of lipids isolated from TX-114 extracts of Δ*aliA* and Δ*aliB* strains. Deletion of *aliA* alone resulted in the complete absence of methylthio-archaeol, while deletion of *aliB* did not reduce the level of thioether-linked archaeol modification. This result correlates well with our observation that deletion of *aliA* reduced the hydrophobicity of substantially more *Hfx. volcanii* lipoproteins than deletion of *aliB* (Fig. 3). Thus, the loss of this lipid modification in the Δ*aliA*/Δ*aliB* strain^43^ is primarily attributable to *aliA* deletion, underscoring the essential role of AliA in mediating this lipid modification (Fig. 5a). Importantly, the absence of thioether-linked archaeol modification in Δ*aliA* was not due to impaired archaeol synthesis, as total lipid profiles were comparable across all strains, including the wild type (Fig. 5b).

**Fig. 5.**
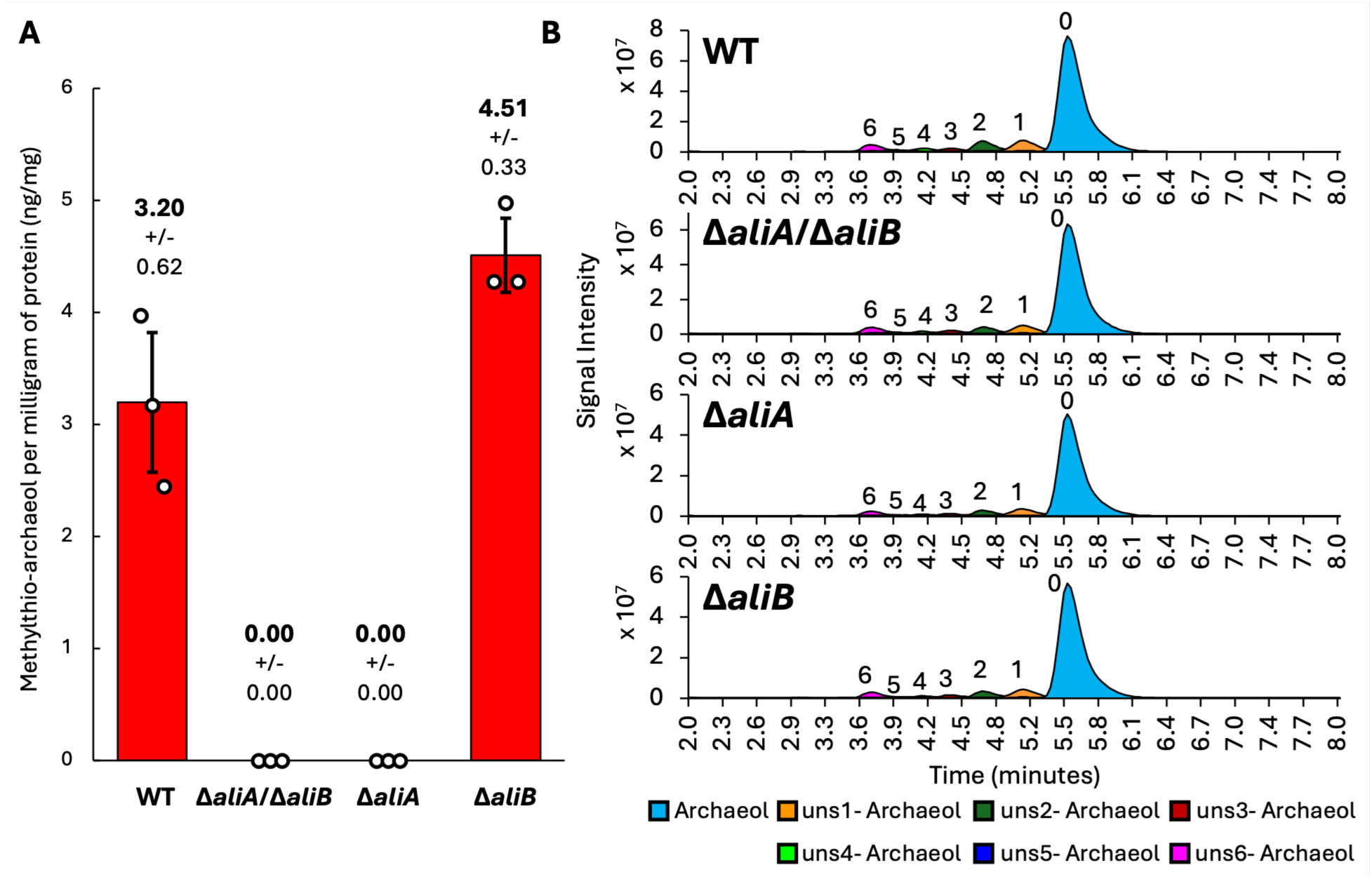
Deletion of *aliA*, rather than *aliB*, led to the complete absence of thioether-linked archaeol in lipoprotein extracts. **(a)** Bar chart showing the average mass of methylthio-archaeol recovered from methyl iodide treated lipoproteins per milligram of starting protein, based on three biological replicates. Error bars represent standard deviations. Statistical significance was assessed using a two-tailed unpaired t-test assuming unequal variances (Welch’s t-test) for comparison between the wild type and Δ*aliB* (p = 0.077). **(b)** As a control, the total lipids of each strain were analyzed. Representative merged extracted ion chromatograms (EICs, m/z values listed in Methods) show comparable relative abundances of saturated and unsaturated diether core lipids (archaeols) in base hydrolyzed biomass from the wild type, Δ*aliA*, Δ*aliB*, and Δ*aliA*/Δ*aliB*.

**Supplementary Fig. 2.**
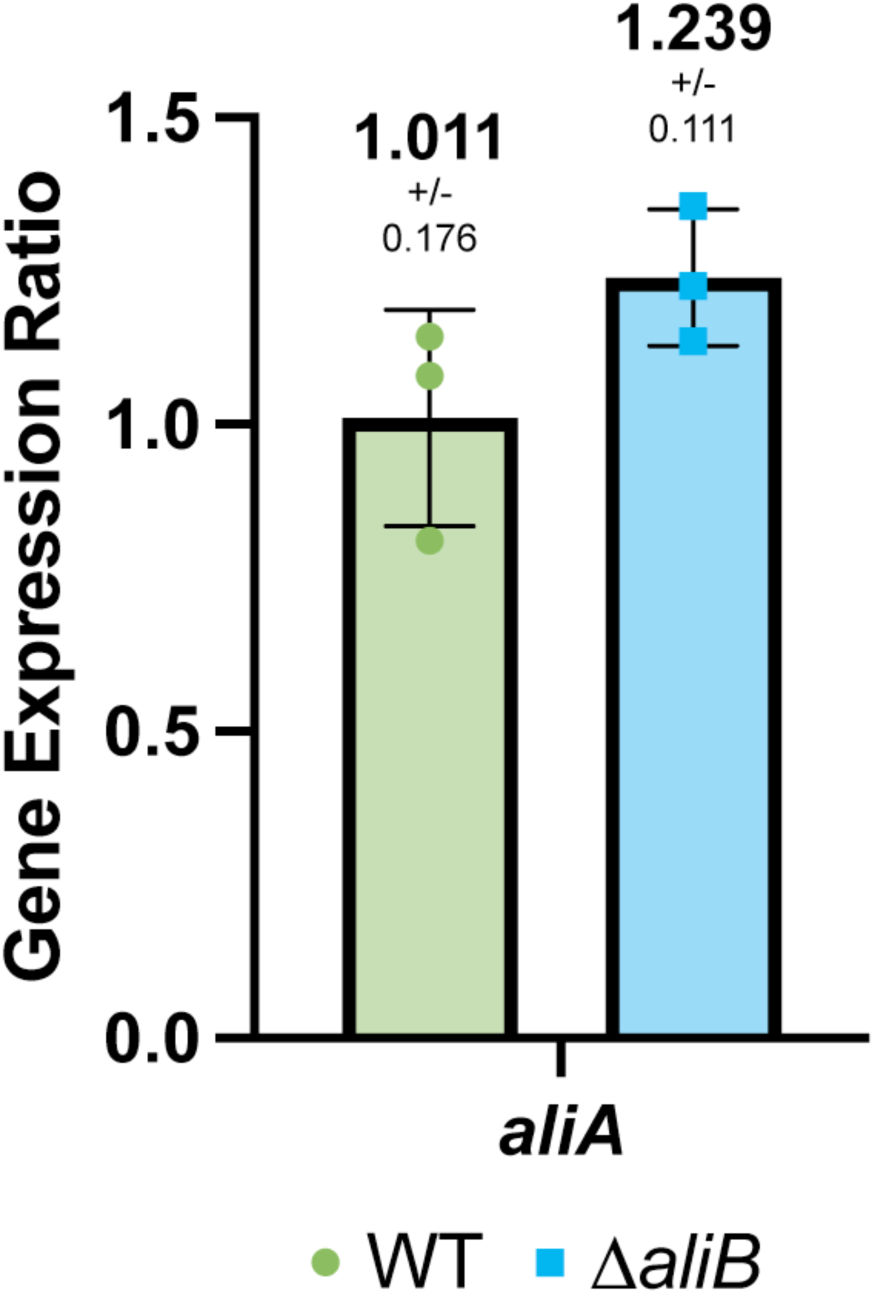
Quantitative PCR showed a slight but non-significant increase in *aliA* expression in Δ*aliB* compared to the wild type (WT) (p-value = 0.174), Bar height represents the mean gene expression ratio normalized to an internal control gene. Error bars indicate standard deviation from three biological replicates. Statistical significance was assessed using a two-tailed unpaired t-test assuming unequal variances (Welch’s t-test) for comparison between the log-transformed gene expression ratios of wild type and Δ*aliB*.

Unexpectedly, while deletion of *aliB* did not reduce the level of thioether-linked archaeol modification, it resulted in a slight increase in methylthio-archaeol levels compared to the wild type (p-value = 0.077, Fig. 5a). To resolve this, we quantified the AliA abundance in the analyzed strains and detected an increased AliA abundance in the absence of AliB. Quantitative PCR (Supplementary Fig. 2, p-value = 0.174) and proteomics data both indicated increased AliA levels in Δ*aliB*. Specifically, the log_2_FC of Δ*aliB* TX phase versus wild-type TX phase was 1.11 (adjusted p-value = 0.07) and 1.60 in the AQ phase (adjusted p-value = 0.04; see Data Availability for more raw data), pointing to potential functional crosstalk between AliA and AliB.

### Downstream effects of *aliA* and *aliB* deletion beyond lipoproteins

Given that lipoproteins participate in diverse cellular processes (Supplementary Data 1), disruption of lipoprotein biogenesis is expected to exert pleiotropic effects extending beyond lipoproteins themselves. Identifying non-lipoproteins whose abundance changes upon *aliA* or *aliB* deletion may therefore provide insight into the molecular basis of the phenotypes observed in Δ*ali* mutants. Moreover, such proteins may represent previously unrecognized components of the archaeal lipoprotein biogenesis pathway. We therefore examined the impact of *aliA* and *aliB* deletion on non-lipoproteins associated with either the TX phase or the AQ phase. TX-associated proteins were defined based on TX vs AQ comparisons within each strain. Proteins exhibiting significant finite log₂ fold enrichment in TX relative to AQ (i.e., detected in both phases but enriched in TX) are shown in Fig. 6a–c (Supplementary Data 3-5). In contrast, proteins detected exclusively in the TX phase—resulting in infinite log₂ fold changes due to missing data in the AQ phase—are shown in Fig. 6d–f (Supplementary Data 6-8). Because these latter comparisons rely on presence/absence behavior and missing values, they are less quantitatively robust than finite enrichment ratios, but they nevertheless highlight strongly phase-specific proteins.

**Fig. 6.**
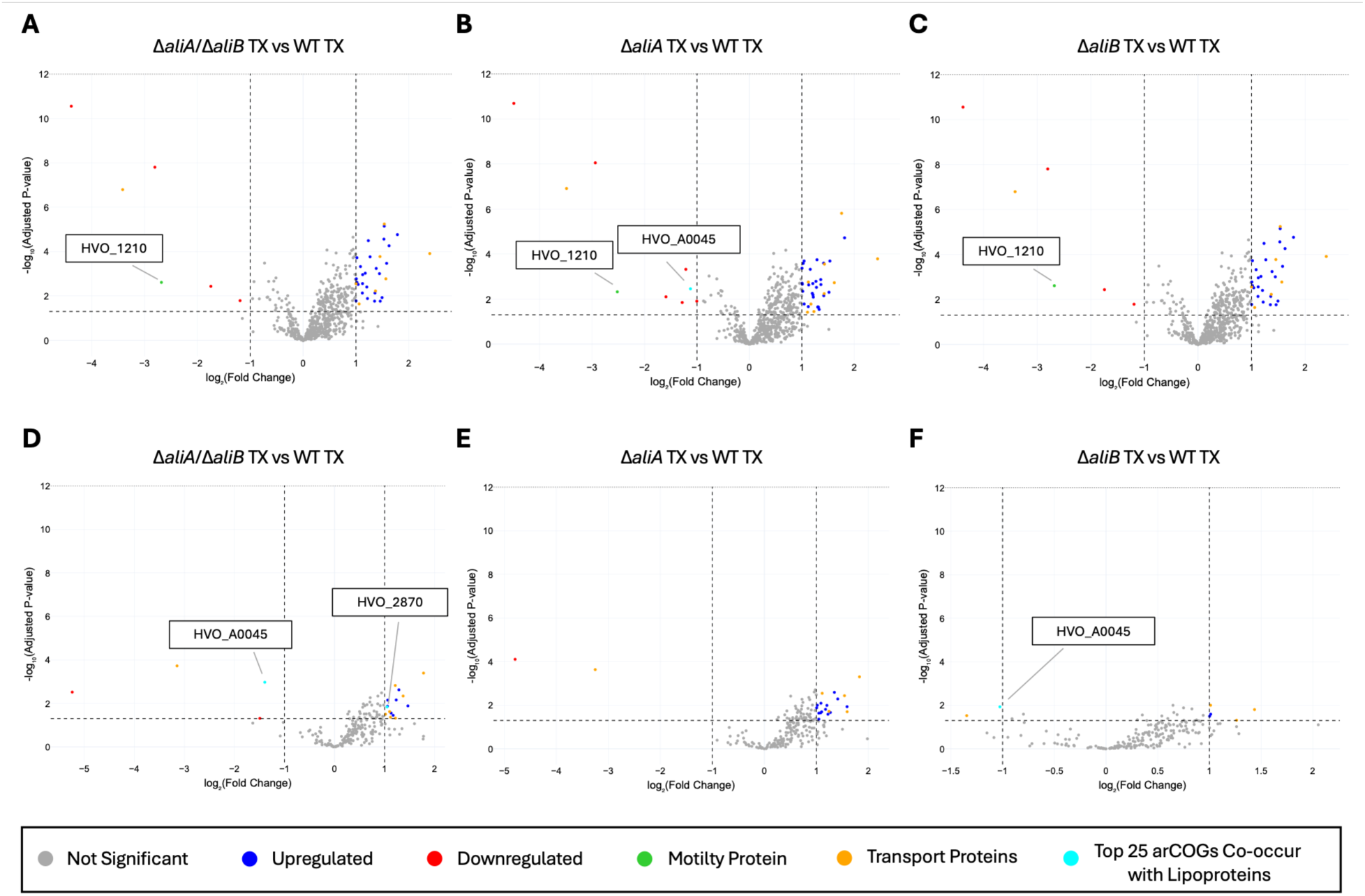
Deletion of *aliA* and *aliB* predominantly perturbs TX-associated non-lipoproteins. Volcano plots show proteins in the TX phase of (a,d) Δ*aliA*/Δ*aliB*, (b,e) Δ*aliA*, and (c,f) Δ*aliB* compared to wild-type *Hfx. volcanii*. Panels a–c display proteins exhibiting significant finite log₂ fold enrichment in TX relative to AQ within each strain (i.e., detected in both fractions but enriched in TX). Panels d–f display proteins detected exclusively in the TX phase of at least one strain, resulting in infinite log₂ fold changes due to absence from the AQ phase. Proteins with |log₂FC| > 1 and adjusted p-value < 0.05 are considered significantly differentially abundant. Selected groups of differentially abundant proteins are highlighted in distinct colors as indicated.

**Supplementary Fig. 3.**
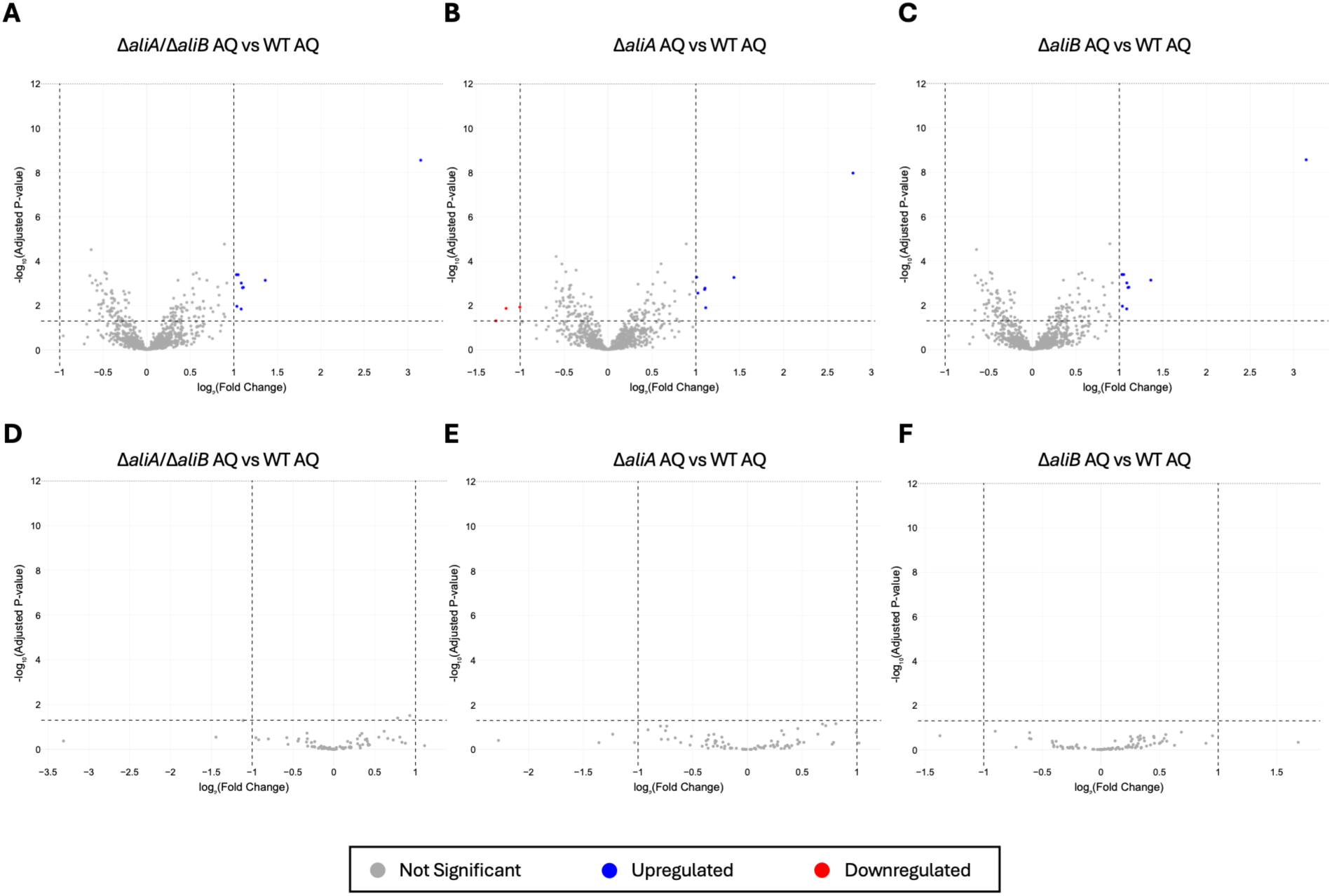
Deletion of *aliA* and *aliB* induces comparatively modest changes in AQ-associated non-lipoproteins. Volcano plots show proteins in the AQ phase of (a,d) Δ*aliA*/Δ*aliB*, (b,e) Δ*aliA*, and (c,f) Δ*aliB* compared to wild-type *Hfx. volcanii*. Panels a–c display AQ-associated proteins exhibiting significant finite log₂ fold enrichment in AQ relative to TX within each strain (i.e., detected in both phases). Panels d–f display proteins detected exclusively in the AQ phase of at least one strain, resulting in negative infinite log₂ fold changes due to absence from the TX phase. Proteins with |log₂FC| > 1 and adjusted p-value < 0.05 are considered significantly differentially abundant. No significantly differentially abundant proteins were detected in panels d-f. Selected functional groups of differentially abundant proteins are highlighted in distinct colors as indicated.

Across all comparisons, a substantial set of non-lipoproteins exhibited significant abundance differences between wild-type and mutant *Hfx. volcanii* strains (Fig. 6, Supplementary Fig. 3, Supplementary Data 3-9). The most pronounced differences were observed in the TX phase (Fig. 6, Supplementary Data 3-8), whereas the AQ phase displayed comparatively fewer and generally more moderate changes (Supplementary Fig. 3, Supplementary Data 9). This pattern indicates that disruption of Ali-dependent lipidation primarily perturbs the membrane-associated proteome rather than the soluble protein pool. Among the TX-associated proteins, the archaellin subunit protein ArlA1 (HVO_1210) was enriched in the detergent phase of all strains but showed reduced abundance in the TX phase of the mutants compared to the wild type (Fig. 6a-c), providing a molecular explanation for their motility defects previously reported^43^. In addition, a large number of transport proteins were more abundant in the TX phase of the mutant strains than in the wild type (Fig. 6), likely reflecting a downstream response to impaired lipidation of ABC substrate-binding proteins, many of which are predicted to be lipoproteins (Supplementary Data 1).

Previously, we identified a comprehensive set of arCOGs (Archaeal Clusters of Orthologous Genes)^58,59^ that most frequently co-occur with lipoproteins as promising candidates for archaeal lipoprotein biogenesis components, including arCOG02177 (*aliA*)^43^. We therefore assessed whether the top 25 arCOGs from the list were differentially represented between mutant and wild-type strains. Among these, HVO_A0045 (arCOG01331), encoding an HtpX-like protease, showed reduced abundance in Δ*aliA* when taking only finite log₂ fold changes into account (Fig. 6b). When proteins that were only detected in the TX phase were also considered, decreased abundance of HVO_A0045 was additionally observed in Δ*aliA*/Δ*aliB* and Δ*aliB* strains (Fig. 6d,f). Bacterial HtpX homologs are implicated in the degradation of misfolded envelope proteins and undergo self-cleavage upon membrane perturbation, suggesting that altered abundance of this protease may reflect changes in envelope protein homeostasis following disruption of Ali-dependent lipidation. Furthermore, when proteins only detected in the TX phase were included in the analysis, HVO_2870 (arCOG02642), a predicted autoinducer-2 exporter (AI-2E) family protein, exhibited increased abundance specifically in the Δ*aliA*/Δ*aliB* strain. Members of the AI-2E protein family have been reported to function either in exporting the quorum sensing molecule autoinducer-2^60^ or as Na^+^ (Li^+^)/H^+^ antiporter^61^. Whether the altered abundance of these proteins reflects direct consequences of Ali pathway disruption, indirect effects stemming from impaired lipoprotein maturation and associated cellular stress, or loss of function of individual lipoproteins remains to be resolved. Finally, we analyzed the abundance of ArtA substrates and ArtA itself in the mutants. No significant differences were detected for either, suggesting that the ArtA-dependent prenylation pathway operates largely independent of the AliA/AliB-dependent lipoprotein lipidation pathway in *Hfx. volcanii*.

## DISCUSSION

Protein lipidation occurs across all domains of life, yet the underlying mechanisms in archaea remain poorly understood. Recently, we identified AliA and AliB as the first archaeal enzymes involved in lipoprotein biogenesis, and showed their involvement in archaeal lipoprotein lipidation^43^. Although the two proteins are paralogs, their functions are not redundant. Deletion of *aliA* or *aliB* in *Hfx. volcanii* led to distinct phenotypes in both lipoprotein biogenesis and cell physiology, and cross-complementation did not restore the defects, indicating that the two enzymes possess different enzymatic activities.

In this study, we further explored the functional differences between AliA and AliB by analyzing the lipidation status of predicted lipoproteins in Δ*aliA*, Δ*aliB*, and Δ*aliA*/Δ*aliB*. We employed a large-scale Triton X-114 extraction, which enabled efficient separation of hydrophobic (TX phase, detergent phase) and hydrophilic (AQ phase, aqueous phase) proteins from large culture samples. Subsequent MS of both fractions yielded high proteome coverage, comparable to the most comprehensive single dataset in the Archaeal Proteome Project (Supplementary Fig. 1a,b), underscoring TX-114 extraction as a powerful method for archaeal proteomic analysis, particularly for enhancing the identification of membrane proteins. Overall, TX-114 extraction resulted in the expected separation of hydrophobic and hydrophilic proteins (Fig. 1a,b), supporting its suitability for large-scale archaeal proteomic analysis. At the same time, a subset of predicted lipoproteins and transmembrane proteins showed enrichment in the AQ phase, while some predicted cytoplasmic proteins were detected in the TX phase. Such fractionation patterns likely reflect a combination of imperfect predictions, minor cross-contamination, protein–protein interactions influencing phase partitioning, or lower-than-expected hydrophobicity of specific lipid modifications or signal peptide H-domains.

This approach also enabled the large-scale validation of archaeal lipoprotein predictions for the first time. In total, 51 of 88 predicted lipoproteins quantified across all strains showed reduced abundance in the TX phase - but not in the AQ phase – in at least one mutant (Fig. 3a), verifying their identity as lipoproteins. Three previously experimentally confirmed lipoproteins were included in this group (Supplementary Data 2). In addition, the MS data corroborated earlier experimental findings for the predicted lipoprotein HVO_1808 (Supplementary Data 2), indicating that the lipobox prediction might be false positive and that the protein likely is not a true lipoprotein, further demonstrating the robustness of this validation strategy. Considering that current archaeal lipoprotein prediction program is trained on a very limited number of experimentally verified lipoproteins^62^, the large-scale lipoprotein validation presented here provides a critical dataset for refining and expanding these predictive models. It should be noted, however, that lipoproteins for which failed lipid attachment leads to protein instability are not captured by our analysis, as the applied filtering criteria exclude proteins with strongly reduced overall abundance. Future studies that directly identify lipid-modified peptides by mass spectrometry will further strengthen lipoprotein validation. Consistent with our previous findings that Δ*aliA* exhibited more severe phenotypes in growth, morphology, and motility than Δ*aliB*^43^, deletion of *aliA* affected substantially more lipoproteins than deletion of *aliB* (Fig. 3). Specifically, 47 of 51 validated lipoproteins showed reduced TX phase abundance in Δ*aliA*, whereas only three lipoproteins were affected in Δ*aliB*, confirming the dominant role of AliA in *Hfx. volcanii* lipoprotein lipidation (Fig. 3a, Supplementary Table 3).

Complementing the whole-proteome analysis, we examined a Tat-transported lipoprotein HVO_1705 both as proof of concept and as a model substrate to further dissect the roles of AliA and AliB. In Δ*aliA* and Δ*aliA*/Δ*aliB* strains HVO_1705 abundance was significantly reduced and the protein localized predominantly to the cytoplasmic fraction (Fig. 4a,b), suggesting impaired translocation or clearance of the non-lipidated form from the membrane by an unidentified quality-control system. In contrast, HVO_1705 expressed in Δ*aliB* remained largely membrane-associated, including both the mature form and a presumed lipidated but non-cleaved intermediate (Fig. 4b). These observations indicate that AliB is not essential for membrane localization and lipidation of HVO_1705 and may instead be involved in signal peptide cleavage of lipoproteins. Analysis showed that a Tat-transported transmembrane protein, HVO_0844, was unaffected by the deletion of either *ali* gene (Fig. 4c,d), confirming that these effects are specific to lipoproteins.

Consistent with the functional distinction between AliA and AliB, LC-MS analysis of lipids isolated from TX-114 extracts further confirmed the dominant role of AliA in mediating thioether-linked archaeol modification in *Hfx. volcanii*. The cleavage product of this modification (methylthio-archaeol) was absent in both Δ*aliA* and Δ*aliA*/Δ*aliB* (Fig. 5). In contrast, deletion of *aliB* did not abolish the modification and instead resulted in a slight, though not statistically significant, increase in the levels of thioether-linked archaeol (Fig. 5). This trend is consistent with the modest upregulation of *aliA* expression observed in Δ*aliB* by both qPCR and MS, suggesting potential regulatory crosstalk between AliA and AliB.

Beyond the altered abundance of *aliA* in the Δ*aliB* mutant, we observed differential abundance of multiple non-lipoproteins in *ali* deletion strains (Fig. 6a-c). A considerable number of non-lipoproteins exemplifies differential abundance in *ali* deletion strains, especially in the TX phase (Fig. 6a-c). Among these were HVO_A0045 and HVO_2870, which were previously identified as proteins that most frequently co-occur with archaeal lipoproteins^43^, although how their respective functions relate to the lipoprotein biogenesis pathway remains unclear. For HVO_A0045, an HtpX-like protease, reduced abundance in the mutants may reflect its involvement in downstream processes of the Ali pathway, such as releasing membrane-associated lipoproteins to the extracellular environment or activation of its self-cleavage activity in response to membrane stress. Although membrane instability has not been reported for *ali* deletion strains, increased outer membrane permeability has been observed in an *Escherichia coli* Δ*lgt* strain^63^. In contrast, HVO_2870, a predicted AI-2E family transporter, showed increased abundance in the double knockout strain. Given that AI-2E family proteins have been reported to function as Na⁺ (Li⁺)/H antiporters^61^, this increase may represent a compensatory response aimed at maintaining intracellular pH homeostasis in response to membrane perturbations associated with *ali* deletion. Additional proteins showing differential abundance in the mutants likely reflect broader consequences of Ali pathway disruption beyond lipoproteins themselves, including downstream effects of impaired lipoprotein lipidation. Among these, a large number of transport proteins were affected (Fig. 6a-f). In addition, the archaellin subunit proteins HVO_1210 exhibited reduced abundance in all mutants (Fig. 6a-c), providing a molecular explanation for the motility defects observed in these strains^43^.

In summary, by combining large-scale quantitative proteomics with LC-MS analysis of thioether-linked archaeol modifications in wild-type and *ali* mutant strains, we demonstrate that the vast majority of archaeal lipoprotein lipidation in *Hfx. volcanii* is mediated by AliA, with AliB acting in a partially distinct role, likely downstream of lipidation and potentially linked to signal peptide cleavage. Beyond defining the dominant contribution of AliA, our quantitative proteomic analysis revealed changes in the abundance of multiple non-lipoproteins in *ali* mutants, which may point to candidate components or regulators of the archaeal lipoprotein biogenesis pathway. Importantly, this work substantially expands the number of experimentally validated archaeal lipoproteins from fewer than ten to over fifty, providing a robust experimental framework for refining lipoprotein prediction algorithms. Together, these data offer a valuable community resource for studying protein lipidation across archaea, where the underlying mechanisms have remained largely elusive.

**Supplementary Table 4.**
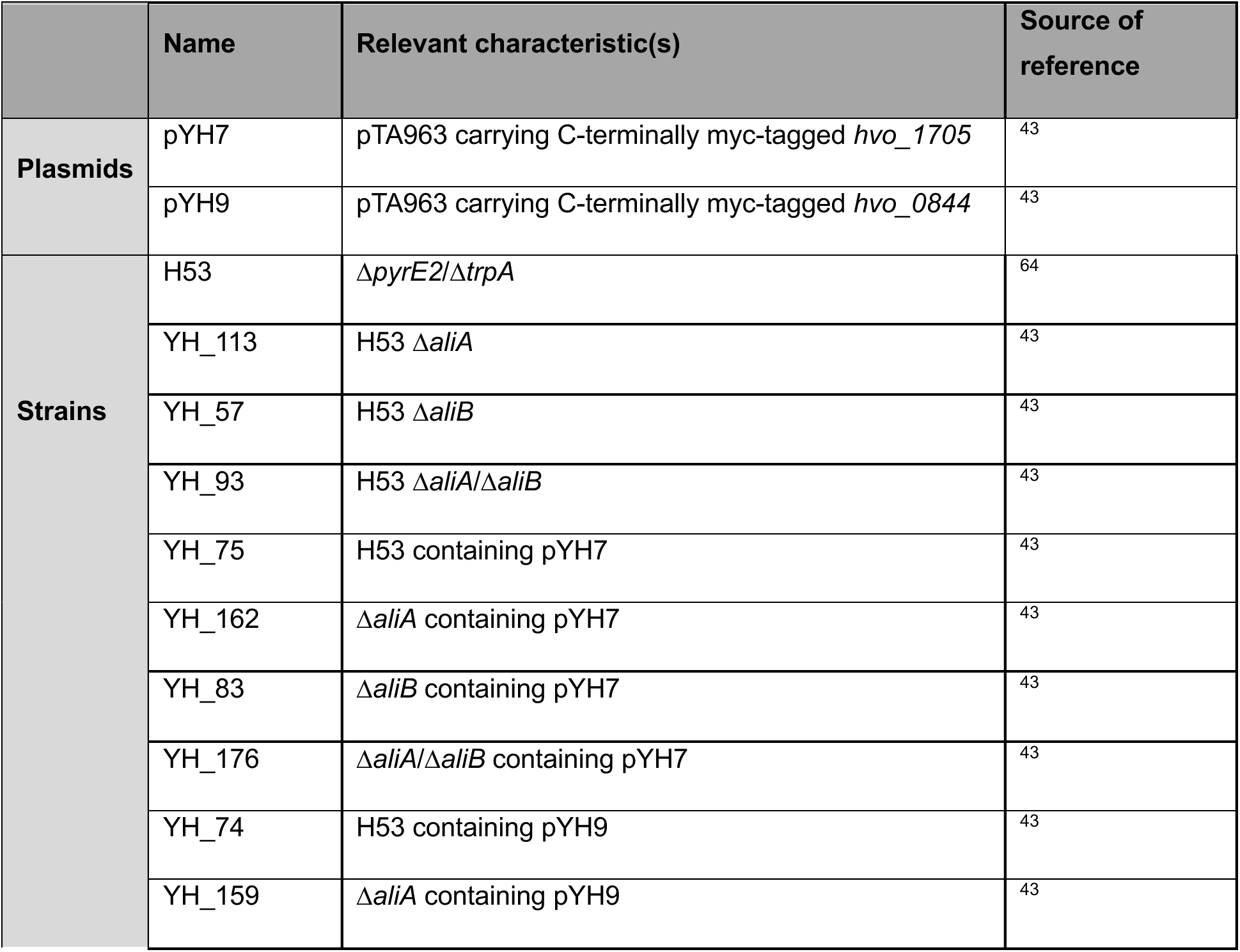
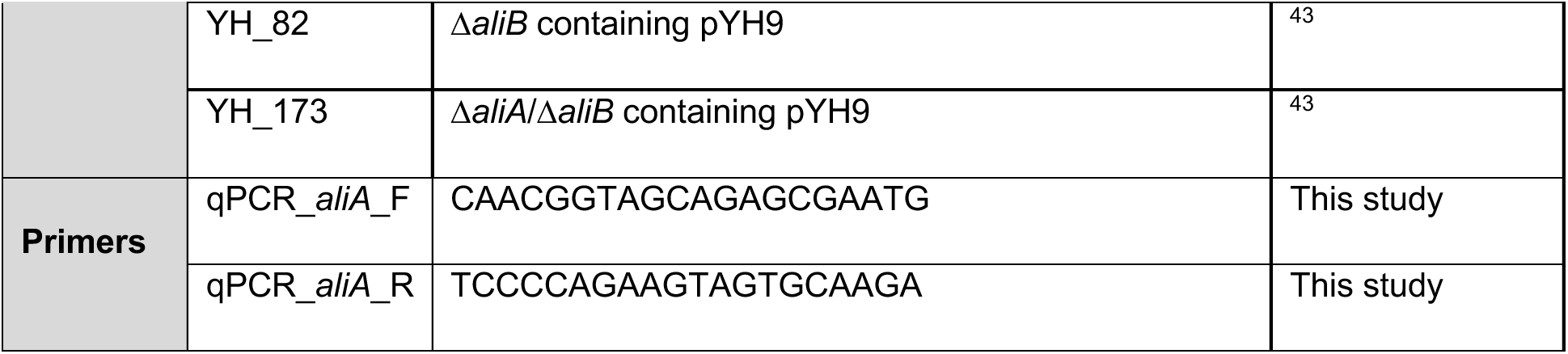
Plasmids, strains, and primers used in this study.

## METHODS

### Growth conditions

Plasmids and strains used in this study are listed in Supplementary Table 4. *Hfx. volcanii* strains were grown at 45 °C in liquid (orbital shaker at 250 rpm with an orbital diameter of 2.54 cm) or on solid agar (containing 1.5% [w/v] agar) semi-defined Hv-Cab medium^65^, supplemented with tryptophan (Fisher Scientific) and uracil (Sigma) at a final concentration of 50 µg ml^-1^ unless otherwise noted. Uracil was left out for strains carrying pTA963-based plasmids.

### Triton X-114 extraction

A single colony of *Hfx. volcanii* was inoculated into 10 ml of liquid medium supplemented with 200 µg ml⁻¹ tryptophan. When the culture reached an OD_600_ of 0.3-0.8, it was used to inoculate 1 l of secondary culture (starting OD_600_ = 0.004) in the same medium. Cells were harvested at OD_600_ = 0.75-0.85 by centrifugation at 7,000 × g for 20 min at 4 °C. The cell pellets were resuspended in 7 ml PBS (pH 7.4) supplemented with 150 µl RQ1 RNase-Free DNase and protease inhibitor, resulting in a final volume of approximately 10 ml. The cell suspensions were transferred to 15 ml Falcon tubes, vortexed thoroughly, and lysed by five freeze-thaw cycles, alternating between freezing in liquid nitrogen for 2 min and thawing at 37 °C for 20 min. Unlysed cells were removed by centrifugation at 9,000 × g for 20 min at 4 °C. Protein concentrations of the clarified lysates were determined using the Pierce™ BCA Protein Assay Kit (Thermo Scientific). Triton X-114 extraction was performed as previously described with slight modifications^43^. Briefly, 1 ml ice-cold 20% Triton X-114 was added to 9 ml of clarified lysate containing the desired amount of protein in a 15 ml Falcon tube. The mixture was gently mixed on a rotator at 4°C for 3 h and then centrifuged at 9,000 × g for 10 min at 4 °C to remove precipitates. The supernatant was transferred to a fresh tube and incubated at 37 °C for 20 min to induce detergent phase separation, followed by centrifugation at 9,000 × g for 10 min at room temperature. The resulting phase separation yielded approximately 9 ml of aqueous (AQ) phase and ∼1 ml of detergent (TX) phase. The AQ phase was transferred to a new 15 ml Falcon tube, and 889 ul of ice-cold 20% TX-114 was added for washing. The TX phase was adjusted to a final volume of 10 ml with PBS for washing. Both samples were incubated on ice for 10 min until clear and then at 37 °C for 20 min until the solution became turbid. Phase-separated samples were centrifuged again at 9,000 × g for 10 min at room temperature. The washed AQ phase was transferred to a new tube and stored at −80 °C for further use. For the washed TX phase, the supernatant was removed, 10 ml methanol was added, and samples were stored at −80 °C overnight to precipitate proteins. Protein precipitates were collected the following day by centrifugation at 10,000 × g for 10 min and the pellet was stored at −80 °C for downstream analysis.

### Mass spectrometry of proteins in TX and AQ phases

Triton X-114 extraction was performed as described above, starting with cell lysates containing 60 mg of total proteins. Protein precipitates from the TX phase were resuspended with 200 µl or 300 µl PBS buffer with protease inhibitor. The protein concentration was measured using the BCA Protein Assay Kit. Mass spectrometry sample preparation was performed as previously described^66^. Briefly, 50 µg of protein from each sample was reduced with DTT, alkylated with iodoacetamide, and digested with trypsin overnight at 37 °C using S-trap mini spin columns (ProtiFi) following manufacturer’s instructions. Resulting peptides were eluted, dried in a vacuum centrifuge, and desalted using C18 stage-tips as described previously^67^. Mass spectrometric analysis was performed, as described in Schiller et al. (2024)^66^, on an Orbitrap Eclipse^TM^ after separation on a nanoEase M/Z Peptide BEH C18 column.

### Proteomics data processing and statistical analysis

Peptide identification and quantification with statistical post-processing followed the same procedures as in Chatterjee et al. (2025)^68^, except that a newer version of the theoretical proteome was used, from March 28^th^, 2023 (https://doi.org/10.5281/zenodo.7794769), containing 4222 proteins. Briefly, peptides were identified using MSFragger, X!Tandem, and MS-GF+ via Ursgal^69^ and quantified through FlashLFQ^70^. Peptide abundances were further processed through MSstats^71^, including log transformation, median normalization, integration on protein level, and calculation of fold-changes (FCs) with corresponding p-values that were adjusted for multiple testing. Log_2_FCs were considered statistically significant if their associated adjusted p-value was ≤0.05.

To compare the distribution of different protein groups in the TX and AQ phases, the whole theoretical proteome was used as an input for the prediction of lipoproteins (comprising lipoproteins processed through the Sec as well as the Tat pathway) through SignalP (v. 6.0)^62^, transmembrane domains through DeepTMHMM (v. 1.0.44)^72^, and type III signal peptides through FlaFind (v. 1.2)^73^. Proteins were assigned to the predicted lipoprotein group, if SignalP results for LIPO(Sec/SPII) or TATLIPO(Tat/SPII) had a confidence of ≥0.9, and if they were not FlaFind positive. The ≥2 TM protein group was defined by DeepTMHMM results that predicted at least two TM regions, as long as these proteins were not FlaFind positive or exhibited SignalP predictions different from “Other”. All proteins that were FlaFind negative and had “Other” as a SignalP prediction, which in combination means that they do not contain any of the secretion signals and had no predicted TM regions, were designated as predicted cytoplasmic proteins.

MSstats results were grouped based on these predicted protein groups, and figures were generated using custom Python scripts (https://doi.org/10.5281/zenodo.18783633), employing the following packages: Pandas^74^, SciPy^75^, Plotly^76^, and Matplotlib^77^. Comparisons between the groups were assessed for statistical significance employing a one-way ANOVA test with Tukey’s post-hoc test, using the SciPy stats package. Protein groups were excluded from analyses downstream of MSstats if all constituent proteins were quantified through proteotypic peptides.

### Lipid processing and analysis

A single colony of *Hfx. volcanii* was inoculated into 5 ml of liquid medium and grown to an OD_600_ of 0.3-0.8. This culture was then used to inoculate 25 ml of fresh medium at a starting OD_600_ = 0.01. Cells were harvested at an OD_600_ = 0.65-0.75 by centrifugation at 7,000 × g for 20 min at 4 °C. Core lipids were obtained from the harvested biomass by base hydrolysis as previously described^43^. For the methyl iodide treatment, lipoproteins were extracted from 80 mg total proteins using Triton X-114. Following extraction, lipoprotein samples were treated with methyl iodide to release thioether bound lipids as previously described^43^.

Core lipids and compounds released from methyl-iodide treatment were analyzed on an Agilent 1260 Infinity II series high-performance liquid chromatography instrument coupled to an Agilent G6125B single quadrupole mass spectrometer. Samples were ionized with electrospray ionization and detected in positive mode with the following parameters: 8.0 l/min drying gas flow rate, 35 psi nebulizer pressure, 3,000 V capillary voltage, and either a 300 °C drying gas temperature (for core lipids) or 250 °C drying gas temperature (for methylthio-archaeol analysis). Core lipids were separated on a Kinetex 1.7 µm XB-C18 100 Å LC column (150 × 2.1 mm), and lipids released from methyl iodide treated lipoproteins were separated on an Agilent Poroshell 120 EC-C18 column (1.9 µm, 2.1 × 150 mm) as described in our previous work^43^. For quantification of the mass of methylthio-archaeol released from each sample, the peak areas of m/z = 683.7, 700.8, and 705.7, (corresponding to the [M+H]+, [M+NH4]+, [M+Na]+ adducts of methylthio-archaeol) were calculated and summed using Agilent MassHunter B.08.00 and inspected manually to confirm the proper integration of extracted ion peaks. The peak area was converted to the mass of methylthio-archaeol injected by the use of a 10-point standard curve as described in our previous work. The methylthio-archaeol (diphytanylglyceryl methyl thioether) standard was synthesized by Pharmaron, Inc. via a synthesis scheme previously described in Hong et al. (2025)^43^.

### Quantitative PCR analysis of *aliA* expression

Three biological replicates of Δ*aliB* and wild type were collected at OD_600_ = 0.8 for each comparison, with 0.5 ml liquid cultures of each strain harvested and pelleted at 4,900 × g for 6 min. RNA was extracted using the Qiagen RNeasy Plus Micro Kit (Qiagen) on the QIAcube Connect automated platform (Qiagen), with lysates homogenized via QIAshredder Columns (Qiagen). RNA extracts were subjected to DNase digestion using the RNase-Free DNase kit (Qiagen) and further purified by RNeasy MinElute Cleanup Kit (Qiagen). RNA concentrations were measured with Qubit RNA Broad Range Assay Kits (Invitrogen), and integrity assessed using RNA ScreenTape on a 4200 Tapestation (Agilent). All samples had RINe values between 8.4-8.8. cDNA was synthesized from 1 µg RNA using the ProtoScript II First Strand cDNA Synthesis Kit (New England BioLabs) with random primers; parallel negative controls without reverse transcriptase (-RT) were prepared. Primers for *Hfx. volcanii* gene *aliA* (Supplementary Table 4) were designed using Primer3^78^ and synthesized as RxnReady™ primers (IDT). Primers for the internal reference gene *eif1a* from prior studies were used^79^. qRT PCR reactions (20 µl) contained 2× iTaq Universal SYBR Green Supermix (Bio-Rad), 5% DMSO, 300 nM primers, and 8 µl of 1:25 diluted cDNA. No-template (NTC) and -RT controls were included; reactions were considered acceptable if ΔCt ≤10 between sample and both controls. Efficiencies for *aliA* and *eif1a* primer sets were determined via standard curves and were 93.0% and 88.7% respectively. Each reaction included three biological replicates, each with three technical replicates, run on a QuantStudio 3 Real-Time PCR System (Applied Biosystems). Thermal cycling conditions were: 95 °C for 2 min, then 40 cycles of 95 °C for 15 s and 60 °C for 30 s. SYBR Green fluorescence (Rox normalized) was recorded at each cycle’s end, and the default melt curve program following the qPCR run verified primer specificity. Relative expression was calculated in Excel using the Pfaffl method^80^, normalizing to *eif1a*, and compared to wild-type controls to determine differential expression.

### Immunoblotting

A single colony was used to inoculate a 5 ml liquid culture. At an OD_600_ of 0.75-0.80, cells from 4 ml liquid culture were harvested by centrifugation at 4,300 x g for 10 min at 4 °C. Cell pellets were then resuspended in 200 µl PBS buffer with 1 mM AEBSF and lysed by freezing (with liquid nitrogen or ethanol with dry ice) and thawing (at 37 °C) four times. 20 µl RQ1 RNase-Free DNase (Promega) was added to cell lysates, and the mixture was incubated at 37 °C for 30 min to degrade DNA. Unlysed cells were removed by centrifugation at 4,300 x g for 10 min at 4 °C, and the supernatant was transferred to a new tube for subsequent fractionation. To fractionate the lysates, 200 µl of cleared supernatant was subjected to ultracentrifugation at 81,000 rpm (231,472 × g) for 30 min at 4 °C using a TLA-100.2 fixed-angle rotor in a Beckman Coulter Optima MAX-TL ultracentrifuge. The resulting supernatant (cytoplasmic fraction) was transferred to a new ultracentrifuge tube, and the pellet (membrane fraction) was resuspended in 200 µl ice-cold PBS with 1mM AEBSF. Both fractions were centrifuged again at 81,000 rpm for 30 min at 4 °C. Pellets were then resuspended in 200 µl PBS with 1mM AEBSF, while the cytoplasmic fraction was transferred to a new tube.

10 µl of HVO_1705 samples and 21 µl of HVO_0844 samples were supplemented with 50 mM dithiothreitol and 1x NuPAGE LDS Sample Buffer. PBS was added to adjust each sample to a final volume of 30 µl. Samples were incubated at 70 °C for 10 min, after which 27 µl of each sample was loaded onto NuPAGE 10%, Bis-Tris mini gels (8 cm x 8 cm, 1.0 mm x 10 wells) with NuPAGE MOPS SDS Running Buffer. After electrophoresis, proteins were transferred to a polyvinylidene difluoride (PVDF) membrane (Millipore) using a semi-dry transfer apparatus (BioRad) at 15 V for 30 min. Subsequently, the membrane was stained with Ponceau S Staining Solution (Cell Signaling Technology) to verify the successful protein transfer. The membrane was then washed for 10 min twice in PBS buffer and blocked for 1 h in 5% non-fat milk (LabScientific) in PBST buffer (PBS with 1% Tween-20). After blocking, the membrane was washed twice in PBST and once in PBS. For detection of the myc tag, anti-Myc antibody (9E10, UPenn Cell Center Service) was diluted 1,000 times with PBS buffer containing 3% bovine serum albumin, which was then used to incubate the membrane overnight at 4 °C. Subsequently, the membrane was washed twice in PBST and once in PBS, followed by a 45-min incubation at room temperature with the secondary antibody solution (Amersham ECL anti-mouse IgG, horseradish peroxidase-linked whole antibody, from sheep (Cytiva) diluted 10,000 times in PBS containing 10% non-fat milk). After incubation, the membrane was washed three times in PBST and once in PBS. HRP activity was assessed using the Amersham ECL Prime Western Blotting Detection Reagent (GE). Membranes were imaged using an Amersham Imager 600, with image analysis conducted using the Amersham Imager 600 Analysis software.

## DATA AVAILABILITY

The LC-MS raw data and quantification for lipid analysis can be accessed through the Stanford Digital Repository [https://purl.stanford.edu/nk164tn8116]. The MS raw files generated for the proteomic analysis in this study have been deposited to the ProteomeXchange Consortium via the PRIDE partner repository with the dataset identifier PXD074718, and all identification and quantification results have been included in the deposited dataset. Raw and processed qPCR data are available at [https://doi.org/10.5281/zenodo.18783633].

All other data supporting the findings of this study are provided within the paper and its supplementary information files. Source data are provided with this paper.

## CODE AVAILABILITY

All scripts for the identification and quantification of peptides and proteins from MS datasets are made available through the Archaeal Proteome Project (http://archaealproteomeproject.org) and its associated GitHub repository (https://github.com/arcpp/ArcPP v1.4.0). Custom Python pipelines used for data filtering and figure generation is available at Zenodo under the identifier DOI: <u>10.5281/zenodo.18783632</u> [https://doi.org/10.5281/zenodo.18783633].

## ACKNOWLEDGEMENTS

We gratefully acknowledge the quantitative proteomics analysis conducted by Dr. Jeremy L. Balsbaugh and Dr. Jennifer C. Liddle of the UConn Proteomics & Metabolomics Facility, a component of the Center for Open Research Resources and Equipment at the University of Connecticut. The authors would like to acknowledge the NIH S10 High-End Instrumentation Award 1S10-OD028445-01A1, which supported this work by providing funds to acquire the Orbitrap Eclipse Tribrid mass spectrometer housed in the University of Connecticut Proteomics & Metabolomics Facility. We also gratefully acknowledge Research Computing at the Rochester Institute of Technology (RIT) for providing computational resources and support that have contributed to the research results reported in this publication. We thank the members of the Proteomics Core Facility of the Max-Planck-Institute of Biochemistry (RRID:SCR_025745) for their assistance with preliminary mass spectrometric analyses of samples containing myc-tagged HVO_1705. We are grateful to Dr. Kira S. Makarova for her insights and guidance on the annotation of specific arCOGs.

Y.H., J.C., and M.P. were supported by the National Science Foundation Grant MBC-2222076 and the University of Pennsylvania Research Fund grant. P.V.W. and A.A.G. were supported by the National Science Foundation Grant EAR-1752564. S.S. gratefully acknowledges support from the College of Science at RIT. S.K. was supported by the Emerson Summer Undergraduate Research Fellowship.

## REFERENCES

1. Jiang, H. et al. Protein Lipidation: Occurrence, Mechanisms, Biological Functions, and Enabling Technologies. Chem. Rev. 118, 919–988 (2018).

2. Banahene, N. et al. Chemical Proteomics Strategies for Analyzing Protein Lipidation Reveal the Bacterial O-Mycoloylome. J. Am. Chem. Soc. 146, 12138–12154 (2024).

3. Braun, V. & Hantke, K. Lipoproteins: structure, function, biosynthesis. in Bacterial Cell Walls and Membranes 39–77 (Springer, 2019).

4. Storf, S. et al. Mutational and bioinformatic analysis of haloarchaeal lipobox-containing proteins. Archaea 2010, 410975 (2010).

5. Ding, W. et al. The Biosynthesis and Applications of Protein Lipidation. Chem. Rev. 124, 12176–12212 (2024).

6. Spang, A. et al. Complex archaea that bridge the gap between prokaryotes and eukaryotes. Nature 521, 173–179 (2015).

7. Zaremba-Niedzwiedzka, K. et al. Asgard archaea illuminate the origin of eukaryotic cellular complexity. Nature 541, 353–358 (2017).

8. Da Cunha, V., Gaïa, M. & Forterre, P. The expanding Asgard archaea and their elusive relationships with Eukarya. mLife 1, 3–12 (2022).

9. Baker, B. J. et al. Diversity, ecology and evolution of Archaea. Nat. Microbiol. 5, 887–900 (2020).

10. Jung, J., Kim, J.-S., Taffner, J., Berg, G. & Ryu, C.-M. Archaea, tiny helpers of land plants. Comput. Struct. Biotechnol. J. 18, 2494–2500 (2020).

11. Pfeifer, K. et al. Archaea biotechnology. Biotechnol. Adv. 47, 107668 (2021).

12. Haque, R. U., Paradisi, F. & Allers, T. *Haloferax volcanii* for biotechnology applications: challenges, current state and perspectives. Appl. Microbiol. Biotechnol. 104, 1371–1382 (2020).

13. Kobayashi, T., Nishizaki, R. & Ikezawa, H. The presence of GPI-linked protein(s) in an archaeobacterium, *Sulfolobus acidocaldarius*, closely related to eukaryotes. Biochim. Biophys. Acta BBA - Gen. Subj. 1334, 1–4 (1997).

14. Kates, M., Palameta, B., Joo, C. N., Kushner, D. J. & Gibbons, N. E. Aliphatic Diether Analogs of Glyceride-Derived Lipids. IV. The Occurrence of Di-O-dihydrophytylglycerol Ether Containing Lipids in Extremely Halophilic Bacteria*. Biochemistry 5, 4092–4099 (1966).

15. Kates, M., Wassef, M. K. & Kushner, D. J. Radioisotopic studies on the biosynthesis of the glyceryl diether lipids of Halobacterium cutirubrum. Can. J. Biochem. 46, 971–977 (1968).

16. Pugh, E. L. & Kates, M. Acylation of proteins of the archaebacteria *Halobacterium cutirubrum* and *Methanobacterium thermoautotrophicum*. Biochim. Biophys. Acta BBA - Biomembr. 1196, 38–44 (1994).

17. Kikuchi, A., Sagami, H. & Ogura, K. Evidence for Covalent Attachment of Diphytanylglyceryl Phosphate to the Cell-surface Glycoprotein of Halobacterium halobium *. J. Biol. Chem. 274, 18011–18016 (1999).

18. Sagami, H., Kikuchi, A., Ogura, K., Fushihara, K. & Nishino, T. Novel Isoprenoid Modified Proteins in Halobacteria. Biochem. Biophys. Res. Commun. 203, 972–978 (1994).

19. Sagami, H., Kikuchi, A. & Ogura, K. A novel type of protein modification by isoprenoid-derived materials. Diphytanylglycerylated proteins in Halobacteria. J. Biol. Chem. 270, 14851–14854 (1995).

20. Abdul Halim, M. F., et al. *Haloferax volcanii* archaeosortase is required for motility, mating, and C-terminal processing of the S-layer glycoprotein. Mol. Microbiol. 88, 1164–1175 (2013).

21. Abdul Halim, M. F., et al. Permuting the PGF Signature Motif Blocks both Archaeosortase-Dependent C-Terminal Cleavage and Prenyl Lipid Attachment for the Haloferax volcanii S-Layer Glycoprotein. J. Bacteriol. 198, 808–815 (2016).

22. Abdul Halim, M. F., et al. Lipid anchoring of archaeosortase substrates and midcell growth in haloarchaea. mBio 11, 10.1128/mbio.00349-20 (2020).

23. Konrad, Z. & Eichler, J. Lipid modification of proteins in Archaea: attachment of a mevalonic acid-based lipid moiety to the surface-layer glycoprotein of Haloferax volcanii follows protein translocation. Biochem. J. 366, 959–964 (2002).

24. Kandiba, L., Guan, Z. & Eichler, J. Lipid modification gives rise to two distinct *Haloferax volcanii* S-layer glycoprotein populations. Biochim. Biophys. Acta BBA - Biomembr. 1828, 938–943 (2013).

25. Siliakus, M. F., van der Oost, J. & Kengen, S. W. M. Adaptations of archaeal and bacterial membranes to variations in temperature, pH and pressure. Extremophiles 21, 651–670 (2017).

26. Gattinger, A., Schloter, M. & Munch, J. C. Phospholipid etherlipid and phospholipid fatty acid fingerprints in selected euryarchaeotal monocultures for taxonomic profiling. FEMS Microbiol. Lett. 213, 133–139 (2002).

27. Lombard, J., López-García, P. & Moreira, D. Phylogenomic investigation of phospholipid synthesis in archaea. Archaea 2012, 630910 (2012).

28. Villanueva, L. et al. Bridging the membrane lipid divide: bacteria of the FCB group superphylum have the potential to synthesize archaeal ether lipids. ISME J. 15, 168–182 (2021).

29. Sinninghe Damsté, J. S., et al. Dominance of mixed ether/ester, intact polar membrane lipids in five species of the order *Rubrobacterales*: Another group of bacteria not obeying the “lipid divide”. Syst. Appl. Microbiol. 46, 126404 (2023).

30. Sahonero-Canavesi, D. X. et al. Disentangling the lipid divide: Identification of key enzymes for the biosynthesis of membrane-spanning and ether lipids in Bacteria. Sci. Adv. 8, eabq8652 (2022).

31. Abdul Halim, M. F., Stoltzfus, J. D., Schulze, S., Hippler, M. & Pohlschroder, M. ArtA-dependent processing of a Tat substrate containing a conserved tripartite structure that is not localized at the C terminus. J. Bacteriol. 199, (2017).

32. Babu, M. M. et al. A database of bacterial lipoproteins (DOLOP) with functional assignments to predicted lipoproteins. J. Bacteriol. 188, 2761–2773 (2006).

33. El Rayes, J., Rodríguez-Alonso, R. & Collet, J.-F. Lipoproteins in Gram-negative bacteria: new insights into their biogenesis, subcellular targeting and functional roles. Curr. Opin. Microbiol. 61, 25–34 (2021).

34. Nguyen, M.-T., Matsuo, M., Niemann, S., Herrmann, M. & Götz, F. Lipoproteins in Gram-positive bacteria: abundance, function, fitness. Front. Microbiol. 11, 582582 (2020).

35. Thompson, B. J. et al. Investigating lipoprotein biogenesis and function in the model Gram-positive bacterium Streptomyces coelicolor. Mol. Microbiol. 77, 943–957 (2010).

36. Gan, K., Gupta, S. D., Sankaran, K., Schmid, M. B. & Wu, H. C. Isolation and characterization of a temperature-sensitive mutant of Salmonella typhimurium defective in prolipoprotein modification. J. Biol. Chem. 268, 16544–16550 (1993).

37. Tokunaga, M., Loranger, J. M. & Wu, H. C. Prolipoprotein modification and processing enzymes in Escherichia coli. J. Biol. Chem. 259, 3825–3830 (1984).

38. Gupta, S. D. & Wu, H. C. Identification and subcellular localization of apolipoprotein *N*-acyltransferase in *Escherichia coli*. FEMS Microbiol. Lett. 78, 37–42 (1991).

39. John H Gardiner, I. V., et al. Lipoprotein *N*-acylation in *Staphylococcus aureus* is catalyzed by a two-component acyl transferase system. mBio 11, e01619 (2020).

40. Armbruster, K. M. & Meredith, T. C. Identification of the lyso-form *N*-acyl intramolecular transferase in low-GC firmicutes. J. Bacteriol. 199, 10.1128/jb.00099-17 (2017).

41. Pohlschroder, M., Pfeiffer, F., Schulze, S. & Halim, M. F. A. Archaeal cell surface biogenesis. FEMS Microbiol. Rev. 42, 694–717 (2018).

42. Giménez, M. I., Dilks, K. & Pohlschröder, M. *Haloferax volcanii* twin-arginine translocation substates include secreted soluble, C-terminally anchored and lipoproteins. Mol. Microbiol. 66, 1597–1606 (2007).

43. Hong, Y. et al. Uncovering the prevalence, key biogenesis enzymes, and biological significance of archaeal lipoproteins. Nat. Commun. 16, 8411 (2025).

44. Mattar, S. et al. The primary structure of halocyanin, an archaeal blue copper protein, predicts a lipid anchor for membrane fixation. J. Biol. Chem. 269, 14939–14945 (1994).

45. Tschumi, A. et al. Functional analyses of mycobacterial lipoprotein diacylglyceryl transferase and comparative secretome analysis of a mycobacterial lgt mutant. J. Bacteriol. 194, 3938–3949 (2012).

46. Dautin, N. et al. Role of the unique, non-essential phosphatidylglycerol::prolipoprotein diacylglyceryl transferase (Lgt) in *Corynebacterium glutamicum*. Microbiology 166, 759–776 (2020).

47. Mao, G. et al. Crystal structure of *E. coli* lipoprotein diacylglyceryl transferase. Nat. Commun. 7, 10198 (2016).

48. Sankaran, K. et al. Roles of histidine-103 and tyrosine-235 in the function of the prolipoprotein diacylglyceryl transferase of *Escherichia coli*. J. Bacteriol. 179, 2944–2948 (1997).

49. González de la Vara, L. E. & Alfaro, B. L. Separation of membrane proteins according to their hydropathy by serial phase partitioning with Triton X-114. Anal. Biochem. 387, 280–286 (2009).

50. Hooper, N. M. & Bashir, A. Glycosyl-phosphatidylinositol-anchored membrane proteins can be distinguished from transmembrane polypeptide-anchored proteins by differential solubilization and temperature-induced phase separation in Triton X-114. Biochem. J. 280, 745–751 (1991).

51. Mathias, R. A. et al. Triton X-114 phase separation in the isolation and purification of mouse liver microsomal membrane proteins. Methods 54, 396–406 (2011).

52. Kittelberger, R., Hansen, M. F., Hilbink, F., de Lisle, G. W. & Cloeckaert, A. Selective extraction of bacterial macromolecules by temperature-induced phase separation in Triton X-114 solution. J. Microbiol. Methods 24, 81–92 (1995).

53. Métivier, A. et al. Triton X-114 phase partitioning for the isolation of a pediocin-like bacteriocin from Carnobacterium divergens. Lett. Appl. Microbiol. 30, 42–46 (2000).

54. Kwok, Y. et al. Rapid isolation and characterization of bacterial lipopeptides using liquid chromatography and mass spectrometry analysis. Proteomics 11, 2620–2627 (2011).

55. Schulze, S. et al. The Archaeal Proteome Project advances knowledge about archaeal cell biology through comprehensive proteomics. Nat. Commun. 11, 3145 (2020).

56. Schulze, S., Pfeiffer, F., Garcia, B. A. & Pohlschroder, M. Comprehensive glycoproteomics shines new light on the complexity and extent of glycosylation in archaea. PLoS Biol. 19, e3001277 (2021).

57. Franco, P. H. C. et al. Proteome characterization of single gene deletion mutants lacking zinc finger µ-proteins in *Haloferax volcanii*. microLife uqag005 (2026) doi:10.1093/femsml/uqag005.

58. Makarova, K. S., Wolf, Y. I. & Koonin, E. V. Archaeal clusters of orthologous genes (arCOGs): an update and application for analysis of shared features between Thermococcales, Methanococcales, and Methanobacteriales. Life 5, 818–840 (2015).

59. Makarova, K. S., Wolf, Y. I. & Koonin, E. V. Towards functional characterization of archaeal genomic dark matter. Biochem. Soc. Trans. 47, 389–398 (2019).

60. Rettner, R. E. & Saier Jr., M. H. The Autoinducer-2 Exporter Superfamily. Microb. Physiol. 18, 195–205 (2010).

61. Dong, P. et al. A UPF0118 family protein with uncharacterized function from the moderate halophile Halobacillus andaensis represents a novel class of Na+(Li+)/H+ antiporter. Sci. Rep. 7, 45936 (2017).

62. Teufel, F. et al. SignalP 6.0 predicts all five types of signal peptides using protein language models. Nat. Biotechnol. 40, 1023–1025 (2022).

63. Diao, J. et al. Inhibition of Escherichia coli Lipoprotein Diacylglyceryl Transferase Is Insensitive to Resistance Caused by Deletion of Braun’s Lipoprotein. J. Bacteriol. 10.1128/JB.00149-21 (2021) doi:10.1128/JB.00149-21.

64. Allers, T., Ngo, H.-P., Mevarech, M. & Lloyd, R. G. Development of additional selectable markers for the halophilic archaeon *Haloferax volcanii* based on the *leuB* and *trpA* genes. Appl. Environ. Microbiol. 70, 943–953 (2004).

65. de Silva, R. T. et al. Improved growth and morphological plasticity of Haloferax volcanii. Microbiology 162, (2021).

66. Schiller, H. et al. Identification of structural and regulatory cell-shape determinants in *Haloferax volcanii*. Nat. Commun. 15, 1414 (2024).

67. Rappsilber, J., Mann, M. & Ishihama, Y. Protocol for micro-purification, enrichment, pre-fractionation and storage of peptides for proteomics using StageTips. Nat. Protoc. 2, 1896–1906 (2007).

68. Chatterjee, P. et al. Quorum sensing mediates morphology and motility transitions in the model archaeon Haloferax volcanii. mBio 16, e00906–25 (2025).

69. Kremer, L. P. M., Leufken, J., Oyunchimeg, P., Schulze, S. & Fufezan, C. Ursgal, Universal Python Module Combining Common Bottom-Up Proteomics Tools for Large-Scale Analysis. J. Proteome Res. 15, 788–794 (2016).

70. Millikin, R. J., Shortreed, M. R., Scalf, M. & Smith, L. M. Fast, Free, and Flexible Peptide and Protein Quantification with FlashLFQ. Methods Mol. Biol. Clifton NJ 2426, 303–313 (2023).

71. Kohler, D. et al. MSstats Version 4.0: Statistical Analyses of Quantitative Mass Spectrometry-Based Proteomic Experiments with Chromatography-Based Quantification at Scale. J. Proteome Res. 22, 1466–1482 (2023).

72. Hallgren, J. et al. DeepTMHMM predicts alpha and beta transmembrane proteins using deep neural networks. 2022.04.08.487609 Preprint at 10.1101/2022.04.08.487609 (2022).

73. Esquivel, R. N., Xu, R. & Pohlschroder, M. Novel Archaeal Adhesion Pilins with a Conserved N Terminus. J. Bacteriol. 195, 3808–3818 (2013).

74. McKinney, W. Data Structures for Statistical Computing in Python. SciPy 2010 10.25080/Majora-92bf1922-00a (2010) doi:10.25080/Majora-92bf1922-00a.

75. Virtanen, P. et al. SciPy 1.0: fundamental algorithms for scientific computing in Python. Nat. Methods 17, 261–272 (2020).

76. Inc, P. T. Collaborative data science. https://plot.ly (2015).

77. Hunter, J. D. Matplotlib: A 2D Graphics Environment. Comput. Sci. Eng. 9, 90–95 (2007).

78. Untergasser, A. et al. Primer3—new capabilities and interfaces. Nucleic Acids Res. 40, e115 (2012).

79. Cote, J. A. et al. A Two-Component Regulatory System Mediates Quorum Sensing–Dependent Morphology and Motility Transitions in the Archaeon Haloferax volcanii. 2025.11.10.687552 Preprint at 10.1101/2025.11.10.687552 (2025).

80. Pfaffl, M. W. A new mathematical model for relative quantification in real-time RT–PCR. Nucleic Acids Res. 29, e45 (2001).

